# Genetic mosaics reveal mechanisms of resistance to VEGF signaling loss during angiogenesis

**DOI:** 10.1101/2025.07.31.667893

**Authors:** Irene Garcia-Gonzalez, Susana F. Rocha, Filipa V. Oliveira, Mariya Lytvyn, Maria S. Sanchez-Muñoz, Aroa Garcia-Cabero, Taija Mäkinen, Rui Benedito

## Abstract

The VEGF ligand and its main receptor VEGFR2 are considered to be essential for endothelial differentiation, proliferation, sprouting and survival. Blood vessels cannot form in growing embryos or tissues when VEGF/VEGFR2 signalling is compromised in all cells. The Anti-VEGF blocking antibody is one of the most widely used antibodies in the clinics, blocking angiogenesis in cancer, wound healing or in ischemic diseases. However, vascular resistance to Anti-VEGF has been reported.

Here we used *iFlpMosaics* and *iSuRe-HadCre* to induce and track genetic mosaics of endothelial cells (ECs) lacking VEGFR2 during the entire embryonic and postnatal development. Surprisingly, *Vegfr2^KO^*ECs adapt, proliferate normally and compete with wild-type cells over time, ultimately forming a substantial portion of the capillary network in most organs. We found that *Vegfr2^KO^* ECs are not able to sprout during the first wave of angiogenesis in most tissues, because it is highly VEGF-dependent and favours the growth and mobilization of wild-type ECs. However, due to their inability to respond to VEGF, over time *Vegfr2^KO^*ECs become dominant in veins, which provide for a long term and continuous source of ECs for subsequent waves of VEGF-independent angiogenesis. Comparative scRNAseq analysis of ECs with acute and long-term loss of VEGFR2, revealed both common and organ-specific molecular mechanisms of adaptation and resistance to VEGF signalling loss. This included significant endothelial venousization and the upregulation of ligands for VEGFR1 and VEGFR3. VEGFR1 only partially compensated for VEGFR2 loss in capillary ECs, whereas VEGFR3 only compensated in arterial ECs. Loss of the three VEGF receptors did not compromise venous growth.

This work changes our understanding of the role of VEGF signaling and its receptors in angiogenesis and reveals mechanisms of adaptation and resistance to their loss.

## Introduction

VEGF is the most important vascular endothelial growth factor, and frequently targeted in the clinics to suppress tumor angiogenesis, metastasis and retina disease ^1^. Resistance to VEGF blocking antibodies arises over time which limits their action ^2–4^. However, resistance to the loss of VEGF signalling has not been reported during physiological vascular development.

Embryos or postnatal tissues lacking the activity of this pathway in all cells fail to develop vessels ^5^. VEGFR2 (encoded by the *Kdr/Flk1* gene in mice) is the most important VEGF receptor and has been shown to be essential for angiogenesis in all vascular development contexts studied so far ^6,7^. Germline mutants for this gene, completely lack blood vessels and die very early during embryonic development ^8^. Studies with mouse embryos chimeras formed by wild-type and Vegfr2 knock-out (*Vegfr2^KO^*) progenitor cells showed that these cannot form ECs ^9^. Conditional deletion of *Vegfr2* in ECs after they differentiate with *Tie2-Cre* ^10^, or in postnatal tissues with *Cdh5-CreERT2*, also strongly impairs angiogenesis ^11^, without any evidence of genetic compensation or resistance.

Despite a large body of literature showing how essential is VEGFR2 for endothelial differentiation, proliferation, migration and survival, previous studies deleting *Vegfr2* in all ECs (*Vegfr2^iDEC^*) during embryonic or postnatal organ development, could only analyse its impact on angiogenesis for a relatively short period of time, given its essential role for vascular and hence tissue development and organ function. The severe impairment of angiogenesis and premature lethality of *Vegfr2^iDEC^* embryos and postnatal animals led to a lack of understanding if ECs are able to adapt to the loss of VEGFR2 over time, or if they are able to make certain organ vascular beds in the complete absence of VEGF/VEGFR2 signalling.

Here we used *iFlpMosaics*^12^ and *iSuRe-HadCre* ^13^, to mosaicly induce and track over long periods of time *Vegfr2^KO^* ECs, allowing us to assess the role of this gene during the entire embryonic and postnatal vascular development. We found that *Vegfr2^KO^* ECs can grow and make new blood vessels over time, being only 2-to-6 fold outcompeted during the entire embryonic and postnatal vascular development of all major organs. *Vegfr2^KO^* ECs expand and outcompete wild-type cells in venous vessels, which provide for a continuous source of these mutant ECs for angiogenesis. Single cell RNAseq (scRNAseq) analysis revealed that *Vegfr2^KO^ ECs* adapt over time by upregulating the expression of venous-specific genes (venousization), and ligands for other VEGF receptors. The additional loss of VEGFR1 further compromises capillary EC survival, proliferation and sprouting, whereas the additional loss of VEGFR3 compromises only the arterialization of *Vegfr2^KO^ ECs*.

This work shows that despite its important role for the first wave of angiogenic growth in embryos and most postnatal tissues, VEGF-VEGFR2 signalling is not essential for many subsequent angiogenic processes, being dispensable for the proliferation of a large fraction of ECs and vascular beds *in vivo*. It is also not needed for the homeostasis of several adult organ vascular beds.

This study changes our understanding of the importance of VEGFR2 for angiogenesis and reveals molecular mechanisms of resistance and adaptation to the prolonged absence of VEGF signalling.

## Results

### *Vegfr2^KO^* ECs can proliferate and compete with wild-type cells

Given the essential role for VEGFR2 during angiogenesis, we predicted that single *Vegfr2^KO^* ECs would be very unfit for angiogenesis and completely outcompeted by wild-type ECs during vascular development. To study how mutant and neighbouring wild-type cells compete for vascular growth, we previously developed *iFlpMosaics*^12^. This technology enables the FlpO-dependent ratiometric induction and tracking over long periods of time of cells expressing MTomato-2A-Cre (knock-out for any floxed gene) or MYFP+ (wild-type). To induce these mosaics we used the *Apln-FlpO* allele, that is expressed in the very first angiogenic sprouting ECs in the embryo at E8.5. In female embryos, the X-chromosome located *Apln-FlpO* allele is mosaicly silenced and expressed only in a fraction of ECs ^14^, generating relatively few mutant and wild-type cells, which is compatible with embryonic development (**Fig. 1a**). Analysis of major organs of *Apln-FlpO iFlpMosaic Vegfr2^Flox/Wt^* (*Vegfr2^HET-E^*^8.5^*^-P7^*) animals at postnatal day 7 (P7) revealed that *Vegfr2^HET^*CD31+ ECs expanded less than wild-type cells in most organs (**Fig. 1b**), showing that a 50% loss of VEGFR2 signalling already impacts these cells ability to expand. Surprisingly however, ECs with full loss of VEGFR2 (*Vegfr2^KO-E8.5-P7^*, Tomato+ in *Apln-FlpO iFlpMosaic Vegfr2^Flox/Flox^* mice) were able to survive and proliferated until postnatal stages, having only a relatively minor competitive disadvantage (2-to-6 fold) when compared with heterozygous or wild-type cells. This competitive disadvantage was more pronounced in the lung endothelium and less in brain ECs (**Fig.1 b and c and Extended Data** Fig. 1a).

**Figure 1:**
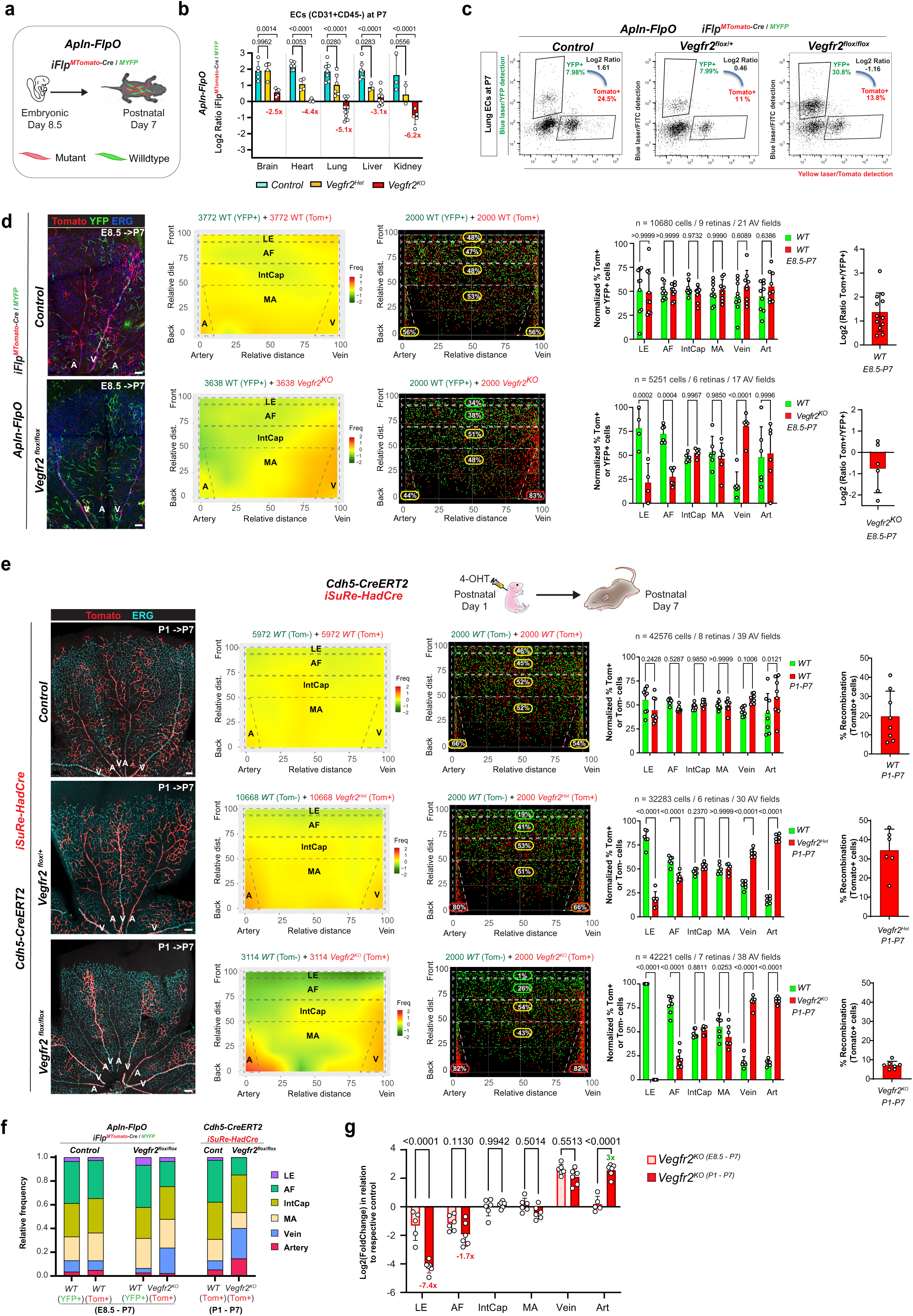
Genetic mosaics reveal how *Vegfr2^KO^* ECs compete with wild-type cells for space. **a,** R26-*iFlp^MTomato-Cre/MYFP^* mosaics were induced with *Apln-FlpO* (recombines endothelial and derived hematopoietic cells at E8.5) on control (wild-type) and *Vegfr2* floxed backgrounds, and different organs were analyzed at P7. **b**, Log2 Ratio of MTomato+/MYFP+ cells detected by FACS of the indicated organs. Absolute fold change is indicated in red. **c,** Representative gating scheme for the ratiometric analysis (see also **Extended Data** Figure 1). **d,** Representative confocal micrographs of control and *Vegfr2^KO-E8.5-P7^* retinas immunostained for Tomato, YFP and ERG (labels EC nuclei). Heatmap and XY dot plot showing the distribution of an equal number of YFP+ and Tomato+ cells (see also **Extended Data** Figure 2). The contribution in percentage of Tomato+ cells to each of the six regions of the retina is indicated both in the XY dot plot and in the chart barplot. Log2 Ratio of Tomato+/YFP+ cells detected in whole mount retina immunostainings, showing an overall competitive disadvantage of Vegfr2^KO^ cells. **e,** Representative confocal micrographs of control, heterozygous (*Vegfr2^HET-P1-P7^*) and homozygous (*Vegfr2^KO-P1-P7^*) deletion of *Vegfr2* retinas induced mosaicly with *Cdh5-CreERT2* and the *iSure-HadCre* reporter at P1, and immunostained for ERG at P7. Heatmap, XY dot plot and charts showing how MTomato+ (Wild-type or *Vegfr2^HET^ or Vegfr2^KO^*) cells distribute in space in relation to MYFP+ (wild-type) cells. **f,** Barplot contrasting the relative frequency of cells in the indicated areas and genotypes. **g,** Log2 of fold change of *Vegfr2^KO-E8.5-P7^* and *Vegfr2^KO-P1-P7^* cells in relation to respective control wild-type cells, showing that *Vegfr2^KO-E8.5-P7^* ECs were more frequent at the leading edge and angiogenic front capillaries than *Vegfr2^KO-P1-P7^* ECs, suggesting adaption overtime. Data are presented as mean values +/-SD. For statistics see Source Data File 1. Scale bars are 100 μm. Abbreviations: LE – leading edge, AF – angiogenic front, IntCap – intermediate capillaries, MA – mature area, A – artery, V-vein.

### *Vegfr2^KO^* ECs can form arteries

It has been previously shown that Vegfr2 activity is essential for Notch activation and the development of arteries ^15–17^. It has also been shown that arteries are formed by tip ECs ^18^, the cells with highest VEGFR2 signalling. Our high resolution spatial zonation data with longterm iFlpMosaics revealed that *Vegfr2^KO-E8.5-P7^* cells were significantly enriched in veins and surrounding capillaries and avoided distal arteries and angiogenic front capillary cells or tip cells in the retina and other organs **(Fig. 1d, Extended Data** Fig.1 **b and Extended Data** Fig. 2 **for heatmapping method**). However, we were surprised to see that a significant fraction of *Vegfr2^KO-E8.5-P7^* ECs were still able to form tip cells and arteries.

To complement these results, and have higher temporal resolution in the analysis, we intercrossed the *Vegfr2* floxed allele with *Cdh5-CreERT2* ^19^ and *iSuRe-HadCre* ^13^ alleles to ensure acute and temporally controlled genetic deletion of *Vegfr2* in all reporter expressing cells. This allowed the mosaic deletion of *Vegfr2* in a subset of ECs at P1. Analysis of retinas at P7 revealed that *Vegfr2^Het-P1-P7^* cells were only excluded from the leading edge of vessels. *Vegfr2^KO-P1-P7^*ECs however were located mainly in veins, surrounding capillaries, and also the more proximal and mature segments of retina arteries (**Fig. 1e and Extended Data** Fig. 2). Notably, ECs with loss of *Vegfr2* from P1 to P7 form the proximal, but not the distal ends, of retina arteries at P7 (**Fig. 1e**). This indicates that when *Vegfr2^KO^* ECs are induced in the initial retina sprouting plexus at P1-P2 and in tip cells, they are able to make arteries forming from P1 to P3. However, given their inability to continue to sprout and form tip cells at the leading edge, they are not able to participate in the subsequent arterial development process occurring from P3 to P7, which is dependent on the incorporation of new tip cells ^20^.

Interestingly, when compared with *Vegfr2^KO-P1-P7^* ECs, *Vegfr2^KO-E8.5-P7^* ECs were much more frequent at the leading edge and angiogenic front capillaries (**Fig.1f, g**) and were still able to make arteries. This data suggests that embryonic *Vegfr2^KO-E8.5-P7^* ECs can adapt to the long-term loss of VEGFR2, and still contribute to venous vascular growth, tip cells and arteries at postnatal stages.

#### *Vegfr2^KO^* ECs proliferate and maintain their long-term lineages in veins

We next determined how *Vegfr2^KO^* ECs can compete with wild-type cells and contribute to growing vascular networks over long periods of time, given the proposed important role of this pathway for EC proliferation ^7,21^. During angiogenesis there are two main locations for EC proliferation, the capillary angiogenic front and veins (**Fig. 2a**). Surprisingly, we found that the surviving *Vegfr2^KO-E8.5-P7^* ECs, proliferate as well as wild-type cells in postnatal brain, heart and lung vessels (**Fig. 2b**).

**Figure 2:**
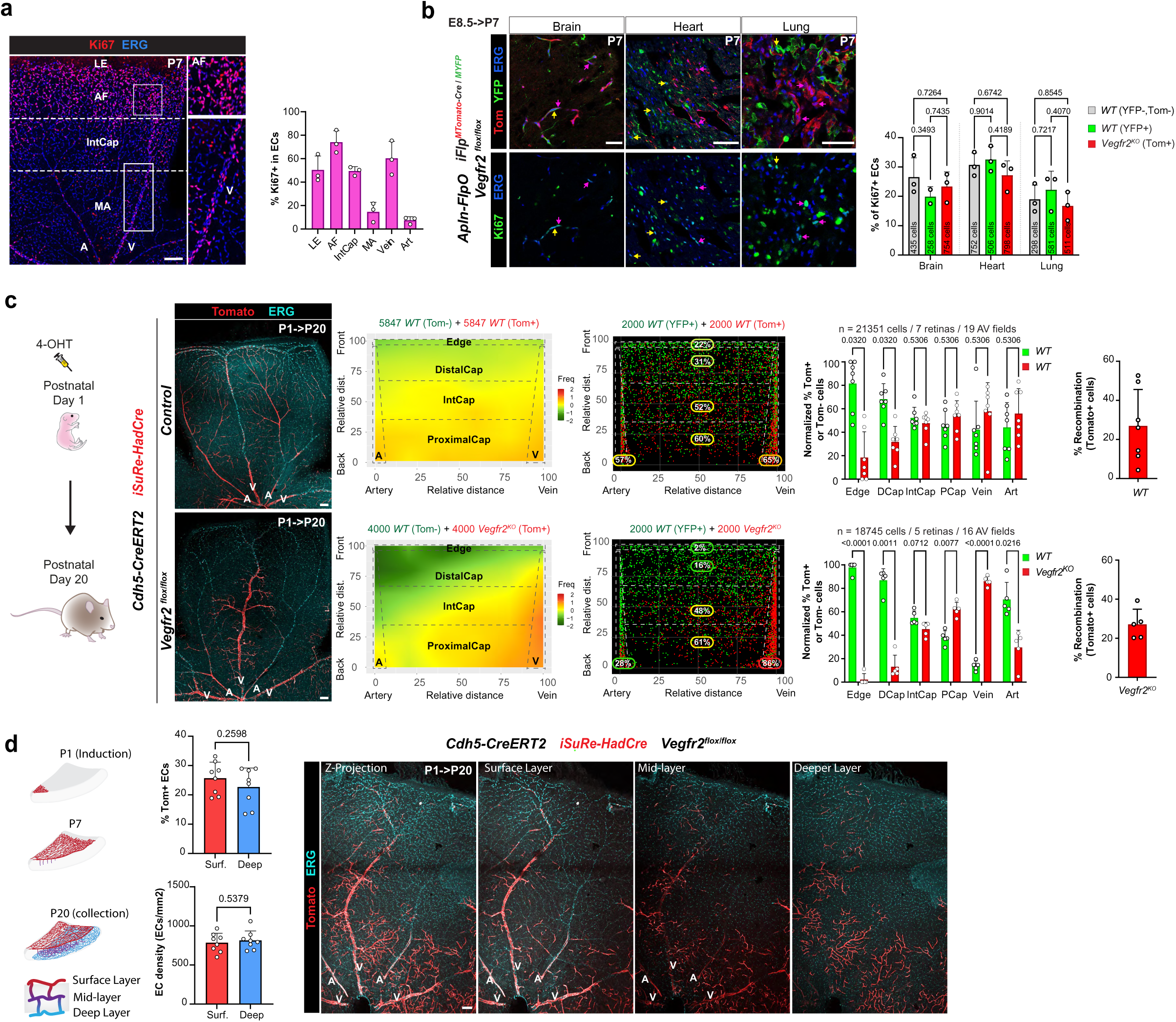
*Vegfr2^KO^* ECs grow and become dominant in veins and their subsequent sprouts. **a,** Representative confocal micrograph of a control P7 retina immunostained for the proliferation marker Ki67 and the EC nuclei marker ERG. Chart showing the frequency of Ki67+/ERG+ (pink) nuclei in the different vascular zones. **b,** Representative confocal micrographs of the indicated organs of *Vegfr2^KO-E8.5-P7^* mice at P7, immunostained for YFP, Tomato, ERG and Ki67. Arrows indicate recombined YFP+ (yellow) and Tomato+ (magenta) ECs that are also Ki67+. Chart with quantifications, showing no difference in the proliferation of *Vegfr2^KO-E8.5-P7^* ECs compared to control cells. **c,** Representative confocal micrographs of control and *Vegfr2^KO-P1-P^*^20^ retinas induced mosaicly with *Cdh5-CreERT2* and the *iSure-HadCre* reporter at P1 and analysed at P20. Heatmap, XY dot plot and chart show that *Vegfr2^KO-P1-P^*^20^ ECs persist long-term in veins and surrounding capillaries. Chart showing degree of recombination at P20. **d,** Scheme depicting the development of the 3 retinal vascular layers over time. Charts with quantification of percentage of Tomato+ ECs and the EC density in the surface and deep vascular layer of the retina, showing that *Vegfr2^KO-P1-P^*^20^ cells contribute to the deeper layers (formed between P7 and P20) at the same percentage they are found in the surface layers (at P7). Data are presented as mean values +/-SD. For statistics see Source Data File 1. Scale bars are 100 μm, except in **b** (50um). Abbreviations: A – artery, V-vein, LE – leading edge, AF – angiogenic front, DCap – Distal capillaries, IntCap – intermediate capillaries, PCap – proximal capillaries.

It has been previously shown that venous ECs provide a continuous source of ECs for angiogenesis, and that venous ECs migrate to capillaries, and from here towards arterial vessels ^18,22,23^. When we induce *Vegfr2^KO^* ECs for longer periods of time, from P1 to P20 (*Vegfr2^IECKO-P1-P^*^20^), we observed that mutant cells stay in the veińs stem where they proliferate and gradually outcompete wild-type cells that move outwards towards the leading edge and adjacent capillaries. The preferential occupation of the venous stem niche by *Vegfr2^KO^* ECs gives them the chance to continue proliferating for longer periods of time, and later form most of the perivenous capillary vessels, during a second wave of VEGF-independent angiogenesis from P7 to P20 (**Fig. 2c**). Venous-derived *Vegfr2^KO^* ECs are able to sprout and make the new retina vessels in the deeper retina layers that form between P7 and P20 (**Fig. 2d**). This agrees with previous findings showing that the tip cells that dive to form the deeper vascular plexus in the retina lack the expression of the VEGFR2 signalling target gene Esm1, and instead their migration is dependent on other pathways, such as Alk/Tgf-beta and Wnt signalling ^24–26^.

Altogether, this data shows that VEGFR2 function is essential for the initial sprouting angiogenesis driven by high VEGF levels in hypoxic embryos and organs, but not for the development of veins, perivenous capillaries and the subsequent steps of capillary angiogenesis and remodelling that occur after the very first vascular plexus is formed in the different organs. Thus, by mobilizing to veins, that are known to provide a continuous source of ECs for angiogenesis ^22^, *Vegfr2^K^*^O^ ECs can continue to proliferate and participate in subsequent steps of vascular development for long periods of time.

#### VEGFR2 withdrawal in angiogenic ECs leads to acute apoptosis followed by adaptation

We previously observed that the distribution of *Vegfr2^KO^*ECs in the retina plexus at P7 was different between animals with long-term (*Vegfr2^KO-E8.5-P7^*) and short-term (*Vegfr2^KO-P1-P7^*) mosaic deletion of Vegfr2. *Vegfr2^KO-E8.5-P7^* ECs were more frequent at the leading edge and angiogenic front than *Vegfr2^IECKO-P1-P7^* ECs (**Fig.1f, g**). Therefore, we hypothesized that *Vegfr2^KO-E8.5-P7^* ECs may get genetically rewired and adapt over time dealing better with the loss of the most important receptor for VEGF. To assess this, we first analysed the frequency of activated ECs and in cycle (Ki67+) of *Vegfr2^KO-E8.5-P7^* ECs having deletion of *Vegfr2* for 17 days, and *Vegfr2^KO-P4-P7^* ECs with deletion of *Vegfr2* for only 3 days. *Vegfr2^KO-P4-P7^* ECs were significantly less proliferative than control cells in the heart and kidney (**Fig. 3a**), whereas the proliferation of *Vegfr2^KO-E8.5-P7^* ECs was similar to control in these organs (**Fig.2b**). This suggests that when deletion of *Vegfr2* is induced embryonically, mutant ECs adapt and later form normal proliferating vessels in these organs. In brain and lungs, even *Vegfr2^KO-P4-P7^* ECs proliferated similarly to control cells (**Fig. 3a**). In the retina, is possible to quantify proliferation in distinct segments of a growing vascular network. Deletion of *Vegfr2* for only 3 days, did not affect retina EC proliferation, even in the VEGF-rich angiogenic front (AF in **Fig. 3b**). However, these *Vegfr2^KO-P4-P7^* mice had a very significant increase in the number of apoptotic (cleaved caspase 3+) ECs (**Fig. 3c**), indicating that during angiogenesis in this organ, VEGFR2 mainly controls endothelial cell migration and survival, but not proliferation. The apoptosis was a reaction to the accute loss of Vegfr2, as *Vegfr2^KO-E8.5-P7^* or even *Vegfr2^IECKO-P1-P7^* did not show any apoptotic ECs (**Fig. 3d**), suggesting adaptation of the surviving cells over time. When the acute loss of *Vegfr2* was induced in all ECs, we observed an immediate depletion of *Vegfr2^KO^* ECs from the VEGF-high angiogenic front, but not from veins and more mature vessels (**Fig. 3e**). Therefore, even if most ECs in the retina plexus upregulate the apoptosis marker in the first days after *Vegfr2* loss (**Fig. 3c**), only the angiogenic ECs that were exposed to VEGF at the time of *Vegfr2* deletion die (**Fig. 3e**).

**Figure 3:**
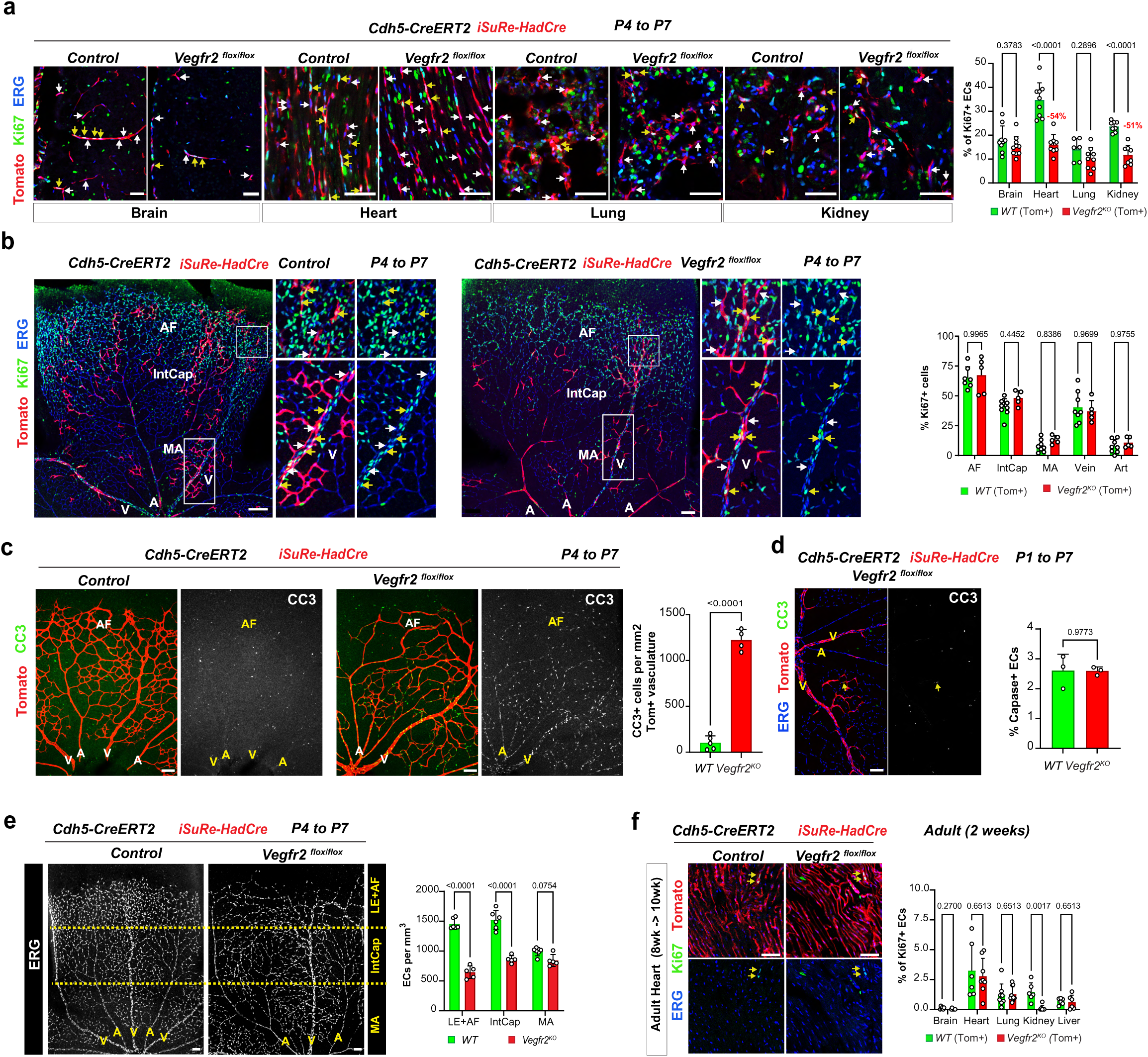
ECs lacking VEGFR2 undergo apoptosis at early stages, but adapt over time. **a,** Representative confocal micrographs the indicated organs of control and *Vegfr2^KO-P4-P7^*mice induced at P4 and analysed at P7. Arrows indicate recombined ECs (Tomato+) that are Ki67+ (yellow) and Ki67-(white), respectively. Chart with the percentage of Tomato+ ECs positive for Ki67+. **b,** Representative confocal micrographs of control and *Vegfr2^KO-P4-P7^* retinas immunostained for Ki67 and ERG. Magnified insets of vein and AF are shown. Arrows indicate recombined ECs (Tomato+) that are Ki67+ (yellow) and Ki67-(white), respectively. Chart with quantifications showing no significant difference in Ki67 positivity between control and *Vegfr2^KO-P4-P7^*ECs. **c,d,** Representative confocal micrographs of retinas immunostained for cleaved caspase 3 (CC3) and with endogenous Tomato signal, and respective quantification chart, showing an increase in EC apoptosis in *Vegfr2^KO-P4-P7^* ECs, after only 3 days of Vegfr2 deletion (from P4 to P7, **c**) but not after a longer period of 6 days (from P1 to P7, **d**). **e,** Representative confocal micrographs of retinas immunostained for ERG and respective quantification chart, showing decreased EC density in the LE+AF and IntCap regions upon *Vegfr2* deletion. **f,** Representative confocal micrographs of adult heart sections of the indicated genotypes and immunostained for Ki67 and ERG and with endogenous Tomato signal. Yellow arrows indicating Ki67+ recombined ECs. Quantification chart showing no difference in EC proliferation, with the exception of the kidney (see also **Extended Data** Figure 3). Data are presented as mean values +/-SD. For statistics see Source Data File 1. Scale bars are 100 μm, except in **a, f** (50 μm). Abbreviations: A – artery, V-vein, AF – angiogenic front, IntCap – intermediate capillaries.

Altogether, our data suggests that ECs lacking VEGF signalling during active angiogenesis suffer from a VEGF signalling addiction and acute withdrawal effect, followed by a period of adaptation to its loss, after which they can form new vessels. ECs more distant from the VEGF-source, and therefore having less VEGF signalling, adapt better to its loss and can expand over time, particularly in veins and perivenous vessels.

To further confirm the insensitivity of more mature ECs to the loss of VEGFR2 signalling, we deleted *Vegfr2* in adult vascular beds, when the levels of VEGF in distinct organs are much lower, and vessels are mostly quiescent. Indeed, only the quiescent liver endothelium showed a strong sensitivity and depletion of ECs after deletion of Vegfr2 for 2 weeks, whereas all other organs maintained their number of ECs (**Extended Data** Fig. 3**, ERG signal**). Ki67/ERG immunostaining in adult quiescent ECs reveals the existence of very few activated and proliferating cells in adult vessels. Loss of VEGFR2 for 2 weeks did not change the Ki67 marker frequency in the adult brain, heart and lung endothelium (**Fig. 3f and Extended Data** Fig. 3), showing their insensitivity to VEGFR2 loss. Only in the kidney and liver, two organs characterized by the possession of fenestrated endothelium, loss of VEGFR2 led to very significant decrease in the frequency of Ki67+ ECs (**Fig. 3f, kidney**), or a significant loss of sinusoidal capillary ECs (**Extended Data** Fig. 3**, liver**).

#### Transcriptional changes after the acute loss of Vegfr2 signalling

Given the observed differences in the biology of *Vegfr2^KO-P4-P7^* and *Vegfr2^KO-E8.5-P7^*ECs, we performed scRNAseq of these cells, to identify possible differences in their transcriptional profiles and potential mechanisms of adaptation to the loss of VEGFR2 signalling in distinct organ vascular beds.

Despite VEGFR2 being one of the most important receptors for angiogenesis, and considered to be essential for EC biology, Vegfr2^KO-P4-P7^ ECs had just relatively minor changes in their transcriptome. The expression of genes related with sprouting/tip cell biology were in general decreased in all organs analysed, whereas the expression of genes related with apoptosis and vascular permeability was increased (**Fig. 4a,b**).

**Figure 4:**
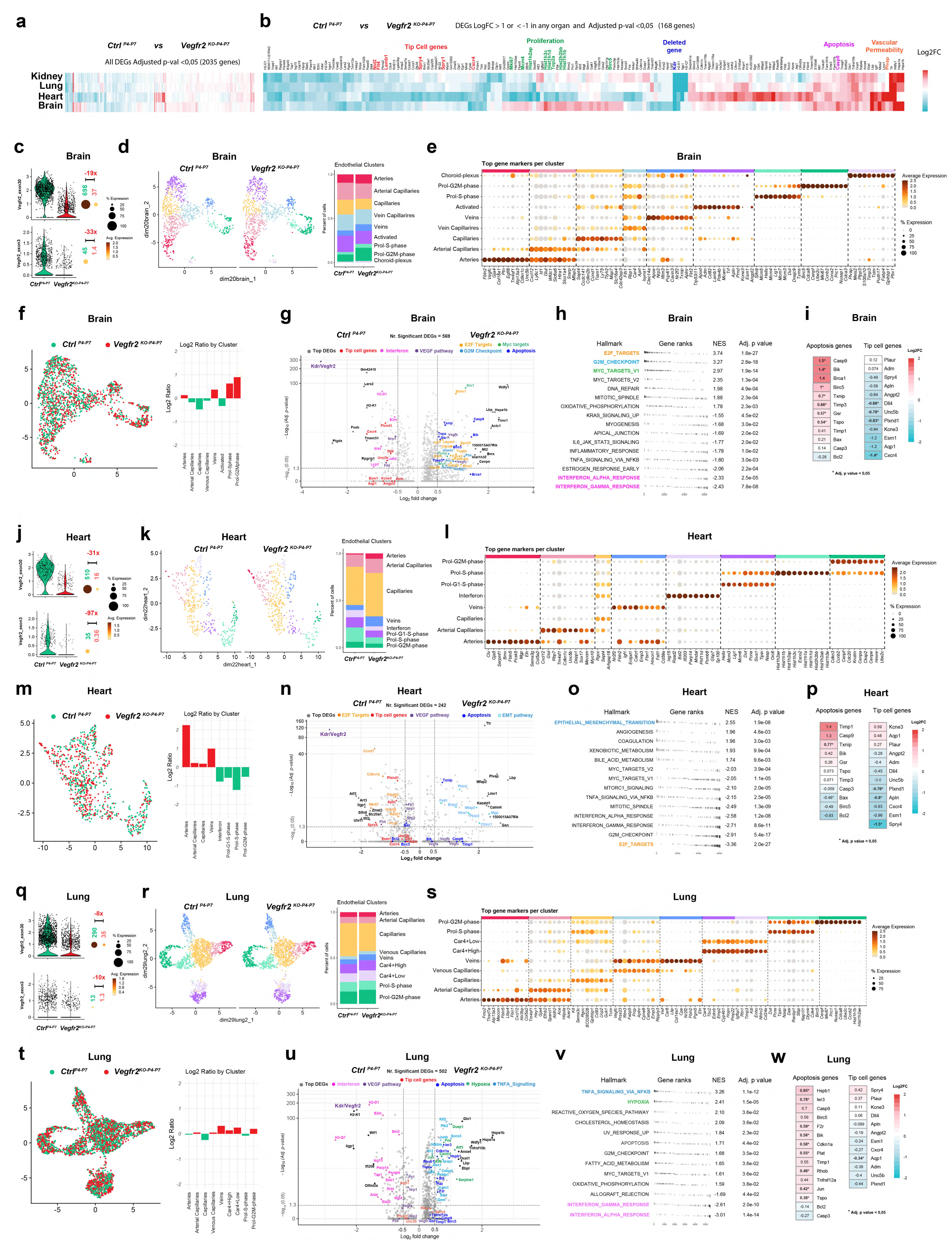
scRNAseq analysis of ECs after the acute loss of VEGFR2 signalling. **a**,**b**, Heatmaps displaying the average logarithmic fold change (Log2FC) of all (**a**) or top (**b**) differentially expressed genes (DEGs) in mutant ECs giving the indicated filter conditions. **c-i,** Analysis of brain ECs. **j-p,** Analysis of heart ECs. **q-w,** Analysis of lung ECs. **c, j, q,** Violin plot and dot plots showing the expression levels of the last exon (exon 30) and the deleted floxed exon (exon 3) demonstrating *Vegfr2* deletion efficiency. Negative numbers in red indicate the fold change between conditions. Green and red numbers above dots represent the average expression multiplied by the percentage of cells expressing the gene to account for absolute expression per group. **d, k, r,** UMAP showing the identified EC clusters and histobar plots showing their proportions per condition. **e, l, s,** Dot plot showing the top cluster marker genes. **f, m, t,** UMAP spatial distribution of control and Vegfr2^KO-P4-P7^ ECs, normalized by cell number. The bar plot shows the log2 ratios of *Vegfr2^KO-P4-P7^* over control cells per EC cluster. **g, n, u,** Volcano plot depicting the top DEGs and also other biologically significant genes when contrasting control and *Vegfr2^KO-P4-P7^* ECs. **h, o, v,** Top significantly up and downregulated hallmark pathways in *Vegfr2^KO-P4-P7^* ECs when compared to control ECs. Highlighted hallmarks match colour code used in (g, n, u). **i, p, w,** Heatmap showing the average Log2FC of apoptosis- and tip cell-related genes in *Vegfr2^KO-P4-P7^* ECs when compared to control ECs.

In the brain, clusters of proliferating ECs, expressing venous capillaries markers, were unexpectedly increased after *Vegfr2* deletion for 3 days (**Fig. 4c-f**). This agrees with the higher clonal expansion of *Vegfr2^KO^*ECs from P1 to P20 in this organ (**Extended Data** Fig. 4a). Clusters comprising capillary and activated/angiogenic ECs (high expression of Apln, Adm, Mcam, Lamb1 and Angpt2) were decreased (**Fig. 4d-f**). In total, only 569 genes were significantly deregulated (**Fig. 4g**). Gene set enrichment analysis (GSEA) revealed a surprising increase in the expression of E2F and Myc target genes, as well G2M, DNA repair and mitotic spindle genes, likely reflecting the increase in EC proliferation (**Fig. 4h and Extended Fig. 4a**). Most hallmarks related with EC interferon and inflammatory response were decreased. The expression of most tip cell marker genes (*Plaur, Kcne3, PlexinD1, Angpt2, Apln* and *Esm1*) was significantly decreased in *Vegfr2^KO-P4-P7^*, while the expression of genes related with apoptosis was in general increased (**Fig. 4i and Extended Data** Fig. 5a-e).

In contrast to the brain, in the heart, *Vegfr2^KO-P4-P7^*capillary ECs were significantly less proliferative (**Fig. 4j,k and Extended Data** Fig. 4b), matching the difference in the Ki67% seen before in sections (**Fig. 3a**). The proportion of mutant ECs found in bigger vessels (arteries and veins) was increased 2 to 4-fold (**Fig. 4k-m**), suggesting a proportional fast depletion of *Vegfr2^KO^* capillary ECs in only 3 days. Opposite to the brain, GSEA of *Vegfr2^KO^*heart ECs revealed the expected decrease in the expression of E2F and Myc target genes, as well a decrease in G2M, mitotic spindle and tip cell genes, which is in accordance with the decrease in proliferation and sprouting of *Vegfr2^KO-P4-P7^* heart ECs (**Fig. 4 n-p**). *Vegfr2^KO^*heart ECs also upregulated genes related with epithelial-to-mesenchymal transition and apoptosis (**Fig. 4n-p**).

In contrast to the two organs shown above, we could not find any significant difference in the proportion of proliferating cells among lung control and *Vegfr2^KO-P4-P7^* ECs (**Fig. 4q-t**), as shown before by immunostaining for Ki67 (**Fig. 3a**). The frequency of the veins cluster was only slightly increased, and the capillary cluster slightly decreased. Despite no significant changes in the clusters proportions, GSEA analysis revealed increased TNFA signalling via NFKB, hypoxia and ROS signalling and a decrease in the interferon response (**Fig. 4u,v**). In this organ there was only a small trend for decreased expression of tip cell marker genes, suggesting the absence of a clear VEGFR2-dependent proliferation and sprouting genetic program in this organ. Several genes related with apoptosis were upregulated (**Fig. 4u and 4w**).

In the kidney, and similar to the heart, *Vegfr2* deletion for 3 days led to an increase in the relative proportion of bigger vessels (arteries and veins) ECs, suggesting a fast depletion of a fraction of capillary ECs in only 3 days (**Extended Data** Fig. 5f-i). The proliferation and clonal expansion of kidney capillary ECs was also significantly reduced (**Extended Data** Fig. 4d **and Extended Data** Fig. 5i), as shown also by immunostaining for Ki67 (**Fig. 3a**). GSEA revealed the expected decrease in the expression of E2F, G2M and mitotic spindle genes and an increase in genes related with p53, xenobiotic metabolism and apoptosis (**Extended Data** Fig. 5j, k), indicative of cellular stress in kidney ECs after short term *Vegfr2* deletion from P4 to P7. Similarly to brain and heart mutant ECs, the expression of tip cell marker genes were in general decreased also in kidney *Vegfr2^KO-P4-P7^*ECs (**Extended Data** Fig. 5l).

Overall, this analysis shows that the acute loss of VEGFR2 induced common but also organ-specific changes in endothelial biology, mainly related with the expression of tip cell genes and cell apoptosis/survival, whereas the impact on EC proliferation was highly variable (**Extended Data** Fig. 5m). We also tried to isolate retina and liver ECs for scRNAseq from the same animals, but capillary ECs in these organs were very strongly depleted after *Vegfr2* deletion from P4 to P7, and therefore sufficient cells could not be obtained.

#### Adaptation to the long-term loss of Vegfr2 signalling through venousization

After the analysis of *Vegfr2^KO-P4-P7^* ECs, we next performed scRNAseq of *Vegfr2^KO-E8.5-P7^*ECs of distinct organs, in order to identify potential differences in gene expression related to the long-term adaptation of cells to the loss of VEGFR2. In contrast to *Vegfr2^KO-P4-P7^*ECs, *Vegfr2^KO-E8.5-P7^* ECs had mostly upregulated, and not downregulated, genes, and importantly a marked upregulation of several venous-enriched genes (**Compare Fig. 4a with 5a)**.

In contrast to brain *Vegfr2^KO-P4-P7^* ECs, brain *Vegfr2^KO-E8.5-P7^*ECs had increased frequencies of capillary cells, particularly venous capillaries (**Fig. 5b-d**), suggesting increased venousization. Similarly to *Vegfr2^KO-P4-P7^*ECs, brain *Vegfr2^KO-E8.5-P7^* ECs had a decreased frequency of activated/sprouting ECs, and lower expression of several tip cell genes (**Fig. 5c-e**). The number of differentially expressed genes, particularly the upregulated genes, was much higher (1621 vs 569 DEGs) in *Vegfr2^KO-E8.5-P7^* ECs, likely as a long term adaptation to the loss of VEGFR2 function (**Compare Fig.5e with 4g**). Whereas in *Vegfr2^KO-P4-P7^* brain ECs genetic pathways related with EC proliferation and cellular activity (E2F targets, Myc, oxidative phosphorylation and glycolysis) were paradoxically increased, in *Vegfr2^KO-E8.5-P7^*ECs they were decreased (**Compare Fig. 4h with Fig. 5f**). This suggests that brain *Vegfr2^KO-E8.5-P7^* ECs adapt to the long-term loss of VEGFR2 signalling by decreasing their cellular activity, and likely in this way their dependence of VEGF signalling for growth and survival. *Vegfr2^KO-E8.5-P7^* ECs also show a strong decrease in the expression of tip cell marker genes (**Fig. 5g**), similar to *Vegfr2^KO-P4-P7^* ECs (**Fig. 4i**).

**Figure 5:**
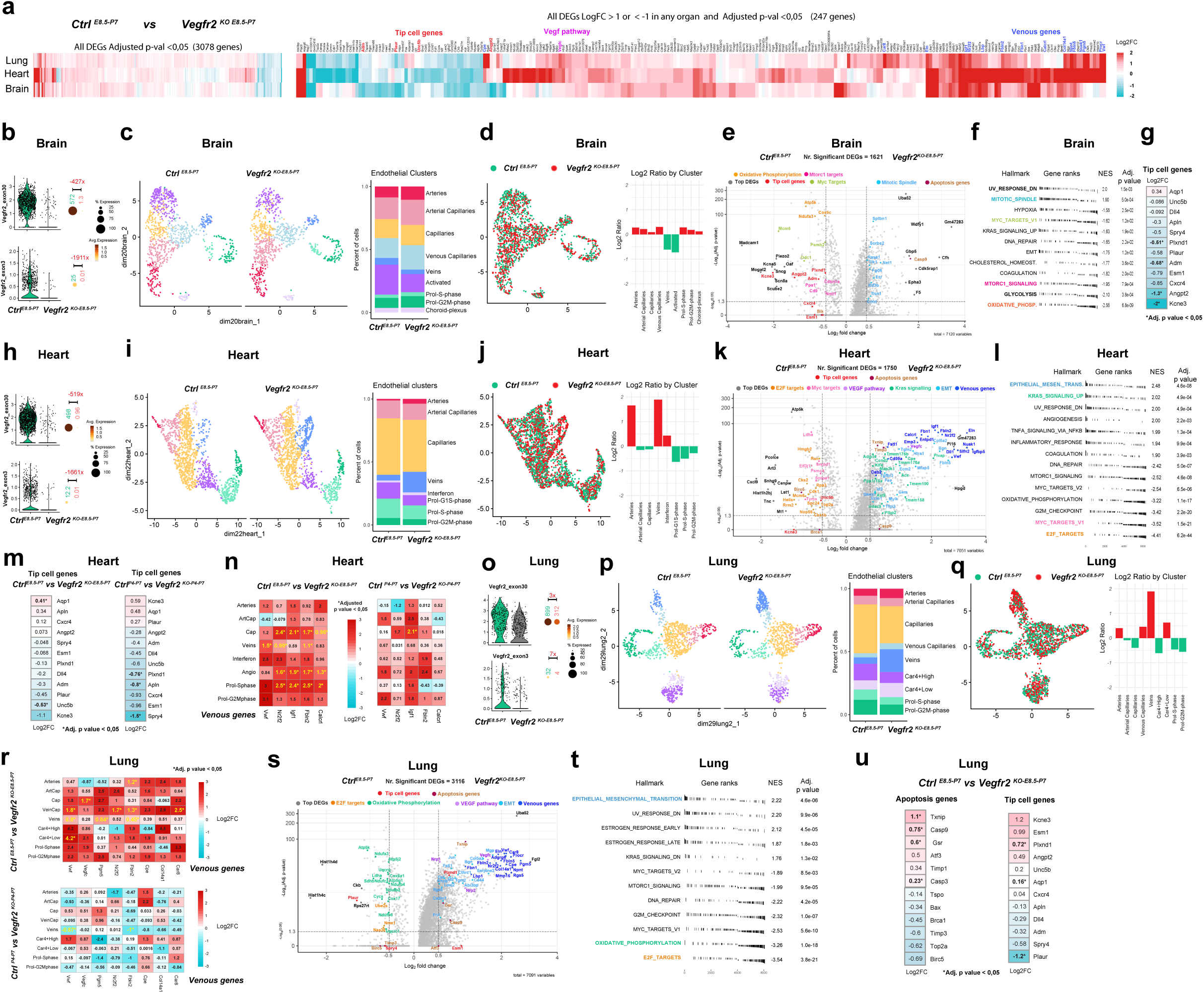
scRNAseq analysis of ECs after long-term deletion of *Vegfr2*. **a**, Heatmaps displaying the average logarithmic fold change (Log2FC) of all or top differentially expressed genes (DEGs) in mutant ECs giving the indicated filter conditions. **b-g,** Analysis of brain ECs. **h-n,** Analysis of heart ECs. **o-u,** Analysis of lung ECs. **b, h, o,** Violin plot and dot plots showing the expression levels of the last exon (exon 30) and the deleted floxed exon (exon 3) demonstrating *Vegfr2* deletion efficiency. Negative numbers in red indicate the fold change between conditions. Green and red numbers above dots represent the average expression multiplied by the percentage of cells expressing the gene to account for absolute expression per group. **c, i, p,** UMAP showing the identified EC clusters and histobar plots showing their proportions per condition. **d, j, q,** UMAP spatial distribution of control and *Vegfr2^KO-P4-P7^* ECs, normalized by cell number. The bar plot shows the log2 ratios of *Vegfr2^KO-P4-P7^* over control cells per EC cluster. **e, k, s,,** Volcano plot depicting the top DEGs and also other biologically significant genes when contrasting control and *Vegfr2^KO-P4-P7^* ECs. **f, l, t,** Top significantly up and downregulated hallmark pathways in *Vegfr2^KO-P4-P7^* ECs when compared to control ECs. Highlighted hallmarks match colour code used in (e, k, s). **g, m, u,** Heatmap showing the average Log2FC of apoptosis- and/or tip cell-related genes in the indicated contrasts. **n, r,** Heatmaps showing the average LogFC of venous genes expression across EC clusters in heart and lung.

In the heart, we confirmed the previous histological analysis showing that indeed heart *Vegfr2^KO-E8.5-P7^* ECs proliferated similar to control cells, and better than *Vegfr2^KO-P4-P7^* ECs (**Compare Fig. 4k,m with Fig. 5i,j**). They also had many more DEGs (1750 DEGs vs 242 DEGs), particularly upregulated (**Compare Fig. 4n with 5k**). GSEA revealed increased epithelial-mesenchymal transition, Kras signalling and angiogenesis, and less expression of metabolic pathway related genes (**Fig. 5k, l**), as seen in the brain. The expression of tip/activated EC genes was not as compromised in *Vegfr2^KO-E8.5-P7^* ECs when compared with *Vegfr2^KO-P4-P7^*ECs (**Fig. 5m**), suggesting some adaptation over time to the loss of VEGFR2. Interestingly, *Vegfr2^KO-E8.5-P7^*ECs were found to be 4 times more frequent in veins (**Fig. 5i, j**), similar to our previous observations in the retina (**Fig. 1d-f**), confirming that the mosaic loss of *Vegfr2* causes mutant ECs to expand more in veins. In comparison with *Vegfr2^KO-P4-P7^* ECs, *Vegfr2^KO-E8.5-P7^*ECs had a strong upregulation of most venous-specific genes, such as Nr2f2, Fbln2, Igf1 and Vwf (**Fig. 5k**). Importantly, this was not only because of the increase in the proportion of vein ECs in these mutants, but also capillary and arterial ECs expressed much more these venous marker genes, indicating a venousization of mutant *Vegfr2^KO-E8.5-P7^* capillary ECs over time (**Fig. 5n**). Endothelial venousization seems to be a key adaptation mechanism to the long-term loss of VEGFR2 signalling, and the reason why they are able to survive, proliferate and compete with wild-type cells over time.

In the lung, the same adaptation and venousization of *Vegfr2^KO-E8.5-P7^*ECs was observed. Mutant cells were 4 times more frequent in veins (**Fig. 5o-q**), and even arterial, capillary or Car4+ ECs significantly upregulated the expression of lung venous genes, like VegfC, Nr2f2, Vwf and Fbln2 (**Fig. 5r**). Similar to the other organs, lung *Vegfr2^KO-E8.5-P7^* ECs also had significantly more DEGs than lung *Vegfr2^KO-P4-P7^* ECs (3116 vs 502), particularly upregulated genes (**Compare Fig. 4u with Fig. 5s**). Volcano and GSEA revealed upregulation of many venous-enriched and epithelial-mesenchymal transition genes, as well as estrogen response genes. Lung *Vegfr2^KO-E8.5-P7^* ECs had less expression of E2F targets and metabolism genes, as seen in the other organs mentioned above (**Fig. 5t**). In contrast to other organs, but similarly to lung *Vegfr2^KO-P4-P7^*ECs, *Vegfr2^KO-E8.5-P7^* ECs did not have a clear deregulation of most apoptosis or tip cell marker genes, suggesting again that in the lung, VEGFR2 does not seem to regulate these processes at P7 (**Fig. 5u**).

Importantly, deletion of *Vegfr2* for 2 weeks in adult organ blood vessels did not induce venousization of adult *Vegfr2^KO^* ECs, it even decreased the expression of venous genes (**Extended Data** Fig. 6), suggesting that venousization is an adaptation mechanism occurring only when angiogenesis is still ongoing.

#### VEGFR1 and VEGFR3 differentially compensate for the absence of VEGFR2

Besides venousization, we have observed a significant upregulation in the expression of other VEGF receptor ligands in *Vegfr2^KO-E8.5-P7^* ECs (**Extended Data** Fig. 7a), such as PGF and VEGFC, that bind and activate VEGFR1 and VEGFR3, respectively ^7^. In the absence of VEGFR2, VEGFR1 is the only other receptor that can bind VEGFA. However, all studies so far showed that VEGFR1 kinase activity is too weak and non-functional ^27,28^, being its extracellular soluble form (sVegfr1/sFlt1) a strong decoy of VEGF ^29^. Therefore, VEGFR1 is considered to mainly antagonize VEGFA/VEGFR2 signalling and angiogenesis, when VEGFR2 is present. However, we have noticed that *Vegfr1* expression was much higher in brain ECs, the organ with less sensitivity to the loss of *Vegfr2* (**Extended Data** Fig. 7b). We also found that during postnatal angiogenesis, sVEGFR1 was 2 to 5 times less expressed than the membrane VEGFR1 (mVEGFR1) form, with exception of brain ECs (**Extended Data** Fig. 7c), suggesting a potential signalling role for mVEGFR1.

To evaluate if VEGFR1 could compensate for the loss of VEGFR2, we induced the mosaic co-deletion of *Vegfr1* and *Vegfr2* in ECs with iSuRe-HadCre from P1 to P7 (*Vegfr1/2^KO-P1-P7^*). The results show that in comparison to *Vegfr2^KO^ ECs*, *Vegfr1/2^KO^*ECs have a further decreased ability to migrate and form retina capillaries, but not veins, where they can still survive and proliferate (**Fig. 6a, 6b**). The ECs surviving deletion of Vegfr1 and Vegfr2 for 6 days, proliferate normally at P7 (**Fig. 6c**). Importantly, this effect is seen when both genes are deleted in single cells, mosaicly, confirming it is cell-autonomous, and not dependent on the loss of soluble Flt1 (sFlt1) expression and function. In contrast to deletion of *Vegfr2*, deletion of both *Vegfr2* and *Vegfr1* impacts the survival rate of the pups (**Fig. 6d**), confirming the requirement for VEGFR1 when VEGFR2 is absent in capillaries. In veins, *Vegfr1/2^KO^* ECs can still survive and clonally expand (**Fig. 6e**).

**Figure 6:**
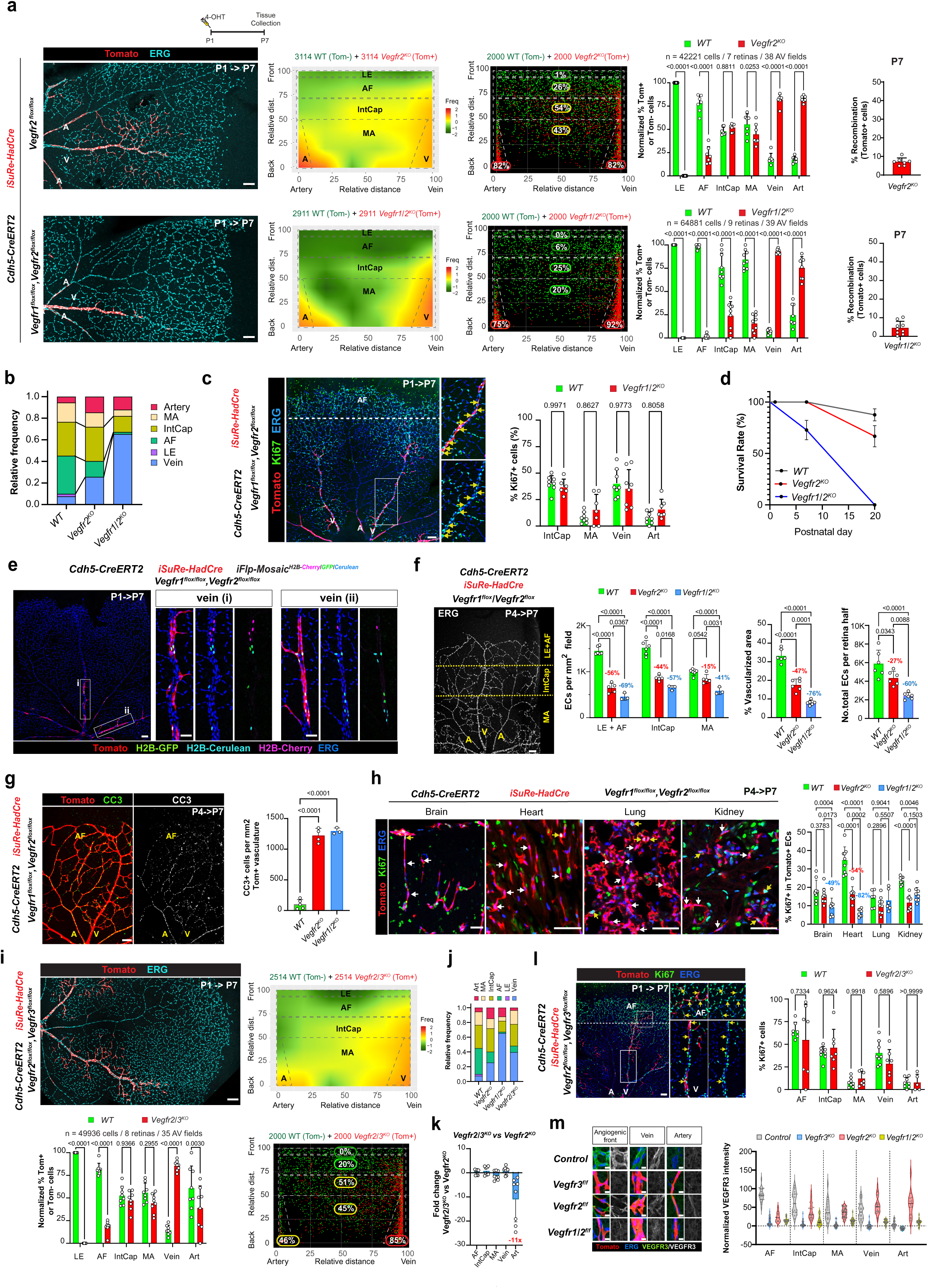
VEGFR1 and VEGFR3 differentially compensate for the absence of VEGFR2 during angiogenesis. **a,** Representative confocal micrographs of retinas vessels of newborn mice induced at P1 and collected at P7. Heatmap, XY dot plot and chart show that *Vegfr1/2^KO^* ECs have a further decreased ability to form retina capillaries, but not veins. Charts on the right show the degree of recombination at P7. **b,** Histobar plot contrasting the relative frequency of Tomato+ cells in the distinct retinal vascular regions at P7, in control, *Vegfr2^KO-P1-P7^* and *Vegfr1/2^KO-P1-P7^* retinas. **c,** Representative confocal micrographs of *Vegfr1/2^KO-P1-P7^* retina immunostained for Ki67 and ERG and with Tomato+/mutant cells. Magnified insets of vein and AF are shown. Yellow arrows indicate Tomato+/Ki67+ cells. Chart with quantifications showing no significant difference in Ki67 positivity between control and *Vegfr1/2^KO-P1-P7^* ECs. **d,** Newborns survival plot showing that double deletion of *Vegfr1* and *Vegfr2* at P1, leads to 100% lethality of pups before P20. **e,** Representative confocal micrographs of *Vegfr1/2^KO-P1-P7^* retina with the additional reporter allele, *iFlp-Mosaic^H2B-Cherry/GFP/Cerulean^*, that allows very sparse labelling of nuclei within the Tomato+/mutant cells. Magnified insets of two veins showing clonal expansion of *Vegfr1/2^KO-P1-P7^* ECs. **f,** Representative confocal micrograph of *Vegfr1/2^KO-P4-P7^* retinas immunostained with ERG and respective quantification charts, showing a further decrease in EC density when compared to *Vegfr2^KO-P4-P7^* retinas. **g,** Representative confocal micrographs of *Vegfr1/2^KO-P4-P7^* retinas immunostained for cleaved caspase 3 (CC3), and respective quantification chart showing a similar increase in EC apoptosis (compare with Figure 3c). **h,** Representative confocal micrographs and quantifications of the % of Ki67+ ECs in the indicated organs of *Vegfr1/2^KO-P4-P7^* mice. Arrows indicate Tomato+/Ki67+ cells (yellow) or Tomato+/Ki67-(white). **i,** Representative confocal micrograph of retinas with double deletion of *Vegfr2* and *Vegfr3* (*Vegfr2/3^KO-P1-P7^*) in ECs from P1 to P7. Heatmap, XY dot plot and chart show that double *Vegfr2/3^KO^*ECs have a further decreased ability to form tip cells and arteries. **j,** Histobar plot contrasting the relative frequency of Tomato+ cells in the distinct retinal vascular regions at P7. Note the significant decrease of arterial cells in *Vegfr2/3^KO-P1-P7^* retinas. **k,** Chart with fold change in contribution of *Vegfr2/3^KO-P1-P7^* ECs relatively to *Vegfr2^KO-P1-P7^* ECs for each of the vascular regions of the retina. **l,** Representative confocal micrograph of a *Vegfr2/3^KO-P1-P7^* retina immunostained for Ki67 and ERG. Magnified insets of vein and AF are shown. Yellow arrows indicate Tomato+/Ki67+ cells. Chart with quantifications showing no significant difference in Ki67 positivity between control and *Vegfr2/3^KO^*ECs. **m,** High magnification confocal micrographs of Tomato+ ECs in the indicated vessels and retina genotypes after ERG and VEGFR3 immunostaining. Violin plot indicating VEGFR3 immunostaining intensity normalized to background levels. Data are presented as mean values +/- SD. For statistics see Source Data File 1. Scale bars are 100um, except in **h** (50 μm) and **m** (10 μm). Abbreviations: A – artery, V-vein, LE – leading edge, AF – angiogenic front, IntCap – intermediate capillaries, MA – mature area.

To better analyse the immediate response of ECs to the loss of both receptors, we induced the short-term full deletion of both genes from P4 to P7. Comparative analysis showed how VEGFR1 provides a considerable survival role to retina capillary ECs in the absence of VEGFR2 (**Fig. 6f, g**). When deleted mosaicly, VEGFR1 partially compensates for the loss of VEGFR2 function in growing retina, brain and heart ECs (**Fig. 6h**). These results show that in the absence of VEGFR2, the weaker VEGFR1 kinase activity provides for key VEGF-dependent signalling, important for capillary EC survival and migration. On the other hand, venous ECs survival and proliferation can occur in the absence of both VEGFR1 and VEGFR2, suggesting that other VEGFA-independent mechanisms enable venous endothelial survival and expansion.

In addition to VEGFR1 and VEGFR2, ECs express also VEGFR3. Previous studies have shown that *Vegfr3* is strongly expressed by capillary and venous ECs, however this receptor cannot bind VEGF (VEGFA), only VEGFC/D, that are key molecules for lymphangiogenesis, but not as relevant for blood vessel angiogenesis ^30^. However, *Vegfr2^KO-E8.5-P7^*ECs showed a very strong upregulation of VEGFC (**Extended Data** Fig. 7a), which could signal to VEGFR3 in the absence of VEGFR2. Paradoxically, loss of VEGFR3 in growing retina ECs was shown to induce angiogenesis, being suggested it negatively regulates VEGFR2 signalling ^31^. When compared with *Vegfr2^iECKO-P1-P7^* ECs, *Vegfr2/3^iECKO-P1-P7^* ECs had a very similar zonal distribution in capillaries and veins, but had a 11-fold lower frequency in arteries (**Fig.6i-k**), suggesting an important role for VEGFC/VEGFR3 in arterial ECs when VEGFR2 signalling is absent. Arterial SMCs are known to express high levels of *Vegfc* ^32–34^ a ligand that can bind both VEGFR2 and VEGFR3. In contrast to arteries, loss of both VEGFR2 and VEGFR3 did not compromise EC proliferation and the development of veins (**Fig. 6l**). We also observed that *Vegfr2^KO^* ECs have decreased VEGFR3 expression only at the angiogenic front, where VEGFA levels are highest, whereas *Vegfr1/2^KO^* ECs have a very strong downregulation of VEGFR3 in all vascular segments, particularly veins and arteries (**Fig. 6m**). Therefore, VEGFR1 is essential to sustain VEGFR3 expression in the absence of VEGFR2 signalling, particularly in mature capillaries, veins and arteries. In the absence of VEGFR2, arterial ECs require VEGFR3 to survive. On the other hand, vein ECs can survive and grow in the absence of all VEGF receptors signalling (**Fig.6c, e, l, m**).

## Discussion

Our study challenges the long-standing view that VEGFR2 signaling is indispensable for endothelial proliferation, migration, and sprouting during angiogenesis ^7,21^. Through the use of *iFlpMosaics* to induce ratiometric genetic mosaics of wild-type and *Vegfr2^KO^*ECs, we demonstrated that these cells are capable of surviving, proliferating, and competing with wild-type ECs over time during the entire embryonic and postnatal organ growth period. Contrary to previous studies suggesting that VEGFR2 is required for all forms of vascular differentiation and development ^9,11,35^, our findings indicate that VEGFR2-independent mechanisms support angiogenesis in certain vascular beds, particularly the long-term development of venous and perivenous capillaries.

We identified significant organ-specific differences in the adaptation to the acute loss of VEGFR2 signalling and significantly extend previous findings ^36,37^. Surprisingly, while angiogenic retina ECs were strongly affected, brain ECs were the least affected, showing no proliferation deficits and retaining angiogenic potential, partially because of VEGFR1 compensation. In contrast, heart, kidney and liver ECs were more sensitive to the acute withdrawal of VEGFR2, with a partial depletion of *Vegfr2^KO^* ECs in capillaries and also reduced proliferation. The lung showed a distinct response, with no major changes in EC proliferation or its clusters, but a shift in transcriptional programs towards inflammation and metabolic adaptations. These findings highlight the heterogeneity of endothelial responses across different vascular beds and suggest that organ-specific factors influence EC resilience to VEGF signaling perturbations.

One of the most striking observations from our study was the long-term adaptation of *Vegfr2^KO^* ECs. While these cells initially faced a competitive disadvantage during early sprouting angiogenesis stages, they ultimately expanded and outcompeted wild-type cells in venous and perivenous regions, indicating that alternative pathways support their survival and proliferation in this niche. Veins are known to provide a continuous source of ECs for capillary angiogenesis and later arterialization ^18,22^, and this venous dominance of *Vegfr2^KO^* ECs, allows them to compete with wild-type cells over time. Our scRNAseq analysis of different vascular beds, further revealed that ECs with prolonged loss of VEGFR2 exhibit a significant shift towards a venous transcriptional signature, marked by upregulation of many venous genes, including *Nr2f2*, a master regulator of veins development ^23,38^. This process, which we term endothelial venousization, appears to be a key adaptation mechanism that enables these cells to persist and contribute to vascular development, despite the full absence of VEGFR2 signalling.

In addition to venousization, our data highlight the compensatory roles of VEGFR1 and VEGFR3 in supporting survival signalling when VEGFR2 is absent in single cells. So far VEGFR1 was mainly found to negatively regulate VEGFR2 signalling during angiogenesis, by being mainly a decoy of the VEGF ligand ^7,21^ . We instead found that VEGFR1 partially compensates for VEGFR2 loss and enables the survival and migration of angiogenic *Vegfr2^KO^* capillary ECs, while VEGFR3 played a similar role in arterial ECs. The ability of these alternative VEGF receptors to sustain EC survival, migration and proliferation at the single cell level, suggests functional plasticity and compensation within the VEGF signaling network, as suggested before by other studies using classical full tissue conditional genetics^11,31,36^.

A major implication of our study is that EC sensitivity to VEGFR2 loss varies depending on the vascular context and the time after its loss. Our experiments revealed that acute loss of VEGFR2 during active angiogenesis resulted in significant EC depletion, particularly of tip cells and other ECs at the angiogenic front, where VEGF signaling is highest. However, *Vegfr2^KO^*ECs in VEGF-low areas and that had undergone long-term adaptation, were able to persist and even contribute to a significant portion of the capillary vessels, particularly those around veins. This suggests that ECs experiencing prolonged VEGFR2 deficiency undergo genetic rewiring in VEGF-low areas, enabling them to survive and grow in the absence of canonical VEGF signaling. The use of ratiometric and trackable iFlpMosaics were key to obtain this finding. This genetic rewiring and long-term adaptation of VEGFR2 mutant cells, may explain why no vascular malformations linked to VEGFR1/VEGFR2 somatic mutations have been reported so far in humans ^39^, despite being the most important VEGF receptors for vascular development ^7,21^.

Our findings also provide insights into resistance mechanisms against anti-VEGF therapies ^40–42^, which are widely used in clinical settings to suppress angiogenesis in cancer and retinal diseases. The ability of ECs to adapt to VEGFR2 loss and engage alternative venous-related angiogenic pathways suggests that targeting VEGFR2 signaling alone may be insufficient for long-term inhibition of pathological angiogenesis. Therapeutic strategies that simultaneously block PLGF-VEGFR1, VEGFC-VEGFR3, or the multiple compensatory and venous-related pathways identified here, may be necessary to overcome resistance to anti-VEGF treatments.

In conclusion, our study reshapes the current understanding of VEGFR2-dependent angiogenesis by demonstrating that ECs possess intrinsic mechanisms to adapt to the long-term absence of VEGFR2 signaling. The discovery of endothelial venousization and compensatory VEGF receptor signaling provides a new framework for investigating vascular development, homeostasis, and therapeutic resistance. Future studies should focus on targeting the identified compensatory molecular pathways underlying these adaptations, in order to overcome anti-VEGF-resistance.

## Acknowledgments

Research in Rui Benedito’s laboratory was supported by the European Research Council (ERC) Consolidator Grant AngioUnrestUHD (101001814), the Ministerio de Ciencia, Innovación y Universidades (PID2020-120252RB-I00 and PID2023-148880OB-I00), and “la Caixa” Banking Foundation (HR22-00316 AngioHeart), all awarded to Rui Benedito. Research in Taija Makinen laboratory was supported by the Knut and Alice Wallenberg Foundation (2018.0218 and 2020.0057) and the Swedish Research Council (2020-02692).

The CNIC is supported by the Instituto de Salud Carlos III (ISCIII), the Ministerio de Ciencia, Innovación y Universidades (MICIU), and the Pro CNIC Foundation. It is recognized as a Severo Ochoa Center of Excellence (grant CEX2020-001041-S funded by MICIU/AEI/10.13039/501100011033).

Microscopy experiments were carried out at the CNIC Microscopy and Dynamic Imaging Unit, an ICTS-ReDib facility co-funded by MCIN (/AEI/10.13039/501100011033) and the ERDF “A way to build Europe” (#ICTS-2018-04-CNIC-16).

We thank the members of the CNIC microscopy, genomics, cytometry, and bioinformatics units for their support throughout the project.

## Authors contribution statement

I.G-G., S.F.R. and R.B. designed most of the experiments, interpreted results, assembled figures, and wrote the manuscript. I.G-G. did confocal microscopy, FACS and scRNAseq bioinformatic analysis, edited text and assembled figures. S.F.R. performed immunostainings, microscopy, FACS analysis, image quantifications in Fiji and graphpad, edited text and figures, and assembled figures. T.M. and F.O. bred mice and provided the samples for the *Vegfr2/Vegfr3* phenotype analysis. M-L, M.S.S-M, A.G.-C, gave general technical assistance with experiments and genotyped the mouse colonies. All authors approved the final version of the manuscript.

## Methods

### Mice

All mouse husbandry and experimentation were conducted using protocols approved by local animal ethics committees and authorities (Comunidad Autónoma de Madrid and Universidad Autónoma de Madrid CAM-PROEX 164.8/20, PROEX 293.1/22and PROEX-288.7/24). The mouse colonies were maintained in racked individual ventilation cages according to current national legislation. The mice had dust and pathogen-free bedding and sufficient nesting and environmental enrichment material for the development of species-specific behavior. All mice had ad libitum access to food and water in environmental conditions of 45–65% relative humidity, temperatures of 21–24 °C and a 12–12 h light– dark cycle. To preserve animal welfare, mouse health was monitored with an animal health surveillance program that followed the Federation of European Laboratory Animal Science Associations (FELASA) recommendations for specific pathogen-free facilities

We used Mus musculus lines mostly on the C57BL6 genetic background, with some having a fraction of 129SV, or B6CBAF1 or DBA genetic backgrounds. Mice were backcrossed to C57Bl6 for several generations. To generate mice for analysis, we intercrossed mice aged between 7 and 30 weeks. We do not anticipate any influence on our data of mouse sex. The following mouse lines were used and intercrossed: *Tg(Cdh5-CreERT2)^1Rha^* ^43^*, Tg(iSuRe-HadCre)* ^13^*, Vegfr2^flox^* ^44^, *Vegfr11^flox^* ^45^, *Vegfr3^flox^*^30^, *Apln-FlpO* ^14^ and the *iFlpMosaic* lines ^12^ : *R26*-*iFlp^MTomato-2A-Cre/MYFP^*, *Tg-iFlp^Chromatin^-Mosaic* and *Tg(INS-CAG-FlpoERT2)*.

To induce CreERT2 or FlpO-ERT2 activity in adult mice, 1g tamoxifen (Sigma-Aldrich, P5648_1G) was dissolved in 50 mL corn oil (stock tamoxifen concentration, 20 mg/mL), and aliquots were stored at - 20C. Animals received intraperitoneal injections of 100 µL of this stock solution (total dose, 2 mg tamoxifen per animal at 80 mg/kg), as indicated in the figures. To activate recombination in pups, 4-Hydroxytamoxifen (4-OHT) was injected at the indicated stages at a dose of 40 mg/kg. All mouse lines and primer sequences required for genotyping are provided in **Supplementary Table 1**.

### Immunostaining of whole retinas

For immunostaining of mouse retinas, eyes were dissected from mouse pups and fixed by incubation with agitation for 20 min in 4% PFA in PBS (diluted from a stock of 16% PFA; EMS 15710). After two washes in PBS, retinas were microdissected from the eyes, 4 incisions made to enable their flat mount later, and refixed in 4% PFA for 45 min with agitation. Dissected retinas were blocked and permeabilized by incubation for 1 h in PBST (0.3% Triton X-100, 3% fetal bovine serum (FBS) and 3% donkey serum in PBS). Samples were then washed twice in PBST and incubated with primary antibodies (**Supplementary Table 2**) diluted in PBST overnight at 4°C with agitation. The next day, retinas were washed 5 × 20 min in PBST diluted 1:2. And after incubated in this same solution for 2 h at room temperature with Alexa-conjugated secondary antibodies (**Supplementary Table 2**). After 3 × 15 min washes in PBS containing 0.15% Triton X-100, retinas were washed 2 × 15 min in PBS and flat mounted in Fluoromount-G (SouthernBiotech). For combining rabbit anti-ERG-647 with other antibodies produced in rabbit, retinas were incubated with 5% rabbit serum for 30 min at room temperature after the Alexa 594 AffiniPure Fab Fragment Donkey Anti-Rabbit secondary antibody, and then incubated with rabbit anti-ERG-647.

### Immunofluorescence on tissue cryosections

For collecting different organs and histological analysis, mice were sacrificed in CO_2_ chambers and 10 ml of PBS were injected into the left ventricles and after perfused with 3.7–4% formaldehyde (ITW Reagents, 252931) at pH 7. The explanted organs were postfixed for 2 h in 4% paraformaldehyde (PFA) (Thermoscientific, 043368.9 M) in phosphate-buffered saline (PBS) at 4 °C with gentle rotation. After three washes in PBS for 10 min each, the organs were stored overnight in 30% sucrose (Sigma) in PBS. The organs were then embedded in optimal cutting temperature (OCT) compound (Sakura) and frozen at −80 °C. Cryosections of 30 µm were cut on a cryostat (Leica), washed three times for 10 min each in PBS and blocked and permeabilized in PBS containing 10% donkey serum (Millipore), 10% fetal bovine serum (FBS) and 1% Triton X-100. The primary antibodies (**Supplementary Table 2**) were diluted in the same buffer and incubated with sections overnight at 4 °C. This step was followed by three 10 min washes in PBS and incubation for 2 h with conjugated secondary antibodies (Supplementary Table 2) and 4,6-diamidino-2-phenylindole (DAPI) in PBS at room temperature. After three washes in PBS, the sections were mounted with Fluoromount-G (SouthernBiotech). For combining rabbit anti-ERG-647 with other antibodies produced in rabbit, retinas were incubated with 5% rabbit serum for 30 min at r/t after the Alexa 594 AffiniPure Fab Fragment Donkey Anti-Rabbit secondary antibody and then incubated with rabbit anti-ERG-647.

### Image acquisition and analysis

For confocal scanning, the immunostained organ sections or whole-mount retinas were imaged at high resolution with a Leica SP8 Navigator confocal microscope fitted with a 10× or 40× objective. Individual fields or tiles of large areas were acquired. All the images shown are representative of the results obtained for each group and experiment. The animals were dissected and processed under exactly the same conditions. Comparisons of phenotypes or signal intensities were made using images obtained with the same laser excitation and confocal scanner detection settings. ImageJ/FIJI v1.54f was used to threshold, select and quantify the objects in confocal micrographs.

For arteriovenous (AV) mapping of ERG (EC nuclei marker) labelled control and mutant endothelial cells (**Extended Data** Fig. 2), a semi-automatic FIJI-script was developed (available at https://github.com/RuiBenedito/Benedito_Lab). Firstly, 6 regions of interest were manually defined by the user: Artery (A), Vein (V), mature capillaries (MA) in between the established A and V, intermediate capillaries (IntCap), angiogenic front (AF) and leading edge (LE). ERG labelling was used to segment endothelial nuclei, classify each nucleus according to the reporter/marker positivity (determined on the percentage of pixel within each nucleus above a given intensity threshold level) and their XY coordinates extracted. After, arterial, venous, distal and proximal AV-boundaries are defined by the user, in order to determine the relative position of each segmented nucleus to each AV field boundaries, and hence allowing the compilation of data extracted from different AV fields into a single XY dot plot or heatmap.

The data collected in Fiji, was then post analysed in RStudio to generate graphs representing the spatial distribution of the cells of interest. For the XY plots and heatmaps, samples were randomly subset to display equal amount of control and mutant cells (MYFP+ versus MTomato+ in the *AplnFlpO, iFlpMosaic* background and MTomato-versus MTomato+ in the *Cdh5-CreERT2, iSuRe-HadCre* background) and in this way make it clear the zonation differences between control and mutant cells. For visualization with XY dot plots, 2000 cells of each group were plotted and for the heatmap generation, the maximal number of cells (determined by the cell type (control or mutant) with less cells) were used. The percentage of each cell type in each AV region was calculated given their spatial distribution per field.

We generated the Fiji macro ‘Script_Reporter_MarkerDetection_Mapping.ijm’ for spatial mapping of ERG (EC nuclei marker) and/or quantification of EC colocalization with reporters (MYFP or MTomato) and other markers (i.e. Ki67) as in Figs. 1,2,3 and 6. R scripts were used for data quantification and heatmapping: (‘Script_Spatial Heatmap’ Figs. 1, 2, 3, 6) and are available at: https://github.com/RuiBenedito/Benedito_Lab. Fiji, Adobe Photoshop CC 24.1.0 and Adobe Illustrator CC v27.1.1 were used for downstream image processing, analysis and illustration.

To quantify the clonal expansion of single endothelial cells expressing the *iSuRe-HadCre* and the *Tg-iFlp^Chromatin^-Mosaic* alleles, images having the MTomato/H2B-Cherry/H2B-EGFP signals were segmented and analysed with the script “Clone_Identification_Quantification”, deposited in https://github.com/RuiBenedito/Benedito_Lab/.

### Flow cytometry and fluorescence-activated cell sorting

Postnatal and adult tissues were dissociated prior to fluorescence-activated cell sorting (FACS). Enzymatic digestion was carried out at 37 °C for 20 minutes using a solution containing 2.5 mg ml⁻¹ collagenase type I (Thermo Fisher), 2.5 mg ml⁻¹ dispase II (Thermo Fisher), and 50 μg ml⁻¹ DNase I (Roche).

Following dissociation, samples were pelleted by centrifugation (450g, 4 °C, 5 min). Pellets were gently resuspended in 1x blood lysis buffer (Biolegend 420301) and incubated on ice for 5 minutes to lyse erythroid cells. After lysis, cells were passed through a 70 μm cell strainer and centrifuged again (450g, 4 °C, 5 min). Cells were resuspended in antibody incubation solution (PBS without Ca²⁺ or Mg²⁺, supplemented with 2% dialyzed FBS; Biowest X0515) together with primary antibodies (APC rat anti-mouse CD31 and APC-Cy7 anti-CD45 – Supplementary Table 2) and HTO-antibodies (Supplementary Table 2) when doing scRNAseq. Samples were then incubated at 4 °C with gentle agitation for 20 minutes.

After labelling, cells were centrifuged (450g, 4 °C, 5 min), washed with antibody incubation solution, centrifuged again, and resuspended in sorting buffer (10% FBS in PBS without Ca²⁺ or Mg²⁺). Stained cells were maintained on ice until acquisition. DAPI (5 mg ml⁻¹) was added immediately prior to FACS analysis.

Cells were analysed using an LSRFortessa cell analyser or sorted using a FACS Aria Cell Sorter (BD Biosciences). For an example of the FACS gating strategy see Extended Fig. 1. BD FACSDiva v8.0.1 and FlowJo v10 were used for data acquisition and analysis.

### Cell isolation for transcriptomic analysis

For RNA-seq analysis, APC rat anti-mouse CD31 (labels ECs), APC-Cy7 rat anti-CD45 (labels blood cells) and HTO-antibodies (BD Biosciences, Supplementary Table 2) were used together to hashtag and distinguish the different organ CD31+ endothelial cells when loaded on the same 10x Genomics port. After antibody incubation, samples were washed twice with antibody incubation solution, centrifuged (450 × *g*, 5 min, 4 °C), and resuspended in sorting buffer (10% FBS in PBS without Ca²⁺ or Mg²⁺). Stained cells were maintained on ice until acquisition. Immediately prior to FACS analysis, DAPI (5 mg ml⁻¹) was added and the cell suspension was passed slowly through a 70 µm cell strainer attached to a 1 ml syringe to remove aggregates.

Live, DAPI-negative, CD31+/CD45- and, endogenous reporter positive or negative cells, were sorted into cold 1.5 ml microcentrifuge tubes containing 300 µl of sorting buffer. Cells were collected using a 100 µm nozzle at 20 PSI under high-purity mode and a flow rate below 3,000 events per second to preserve cell viability and minimize contamination. After sorting, cells were pelleted at 500g for 5 minutes at 4C, and resuspended in 30–40 µl cell-capture buffer (Ca^2+^ and Mg^2+^ free PBS supplemented with 0.04% ultra-pure BSA, Thermofisher AM2616). Viability and cell concentration were determined using a Countess 3 Automated Cell Counter (Thermo Fisher Scientific) before loading the 10x Genomics port.

### Next-generation sequencing sample and library preparation

Single-cell suspensions were loaded onto the 10x genomics Chromium Controller system (10x Genomics) for encapsulation into emulsion droplets. The 10x genomics Chromium single cell 3’ RNA-seq library preparation and sequencing were carried out at the CNIC Genomics Unit according to the manufacturer’s protocols. Each port was loaded with 20 to 50k cells.

Final libraries were sequenced on an Illumina HiSeq 4000 or NextSeq 2000 platforms.

### Single cell RNA data analysis

Single-cell RNA-seq data were aligned and quality controlled by the CNIC Bioinformatics Unit. Clustering, cell type identification and downstream analysis was performed by Irene García-González. The following pipeline was applied for transcript alignment, quantification, quality control, and cell-type classification.

Preprocessing and alignment

Reference transcriptomes were built using the GRCm38 mouse genome and Ensembl gene build v98 (September 2019, sep2019.archive.ensembl.org). Kdr exon 3, Kdr exon 30, Flt1 exon 13b, and Flt1 exon 30 were appended to the reference genome as metadata to enable downstream analysis of gene deletion and the ratio of soluble versus full-length isoforms, respectively. Gene annotations were obtained from the corresponding Ensembl BioMart archive. Raw sequencing reads were aligned and quantified using Cell Ranger v7.1.0.

Cell filtering, demultiplexing and normalization

Downstream analysis was performed using Seurat v4.1.3. Cells were filtered using the following thresholds: >2,000 and <30,000 total counts; >800 and <6000 detected genes; <25% mitochondrial transcript content; <65% of reads in the top 50 genes; <0.1% hemoglobin transcripts; Cdh5>0.1 and Ptprc1>0.1 normalized gene expression.

Cells were demultiplexed by organ using sample-specific hashtag antibody signals (**Supplementary Table 2**), and hashtag-identified doublets were excluded. All subsequent analyses were conducted separately for each organ.

Filtered count data were log-normalized and scaled prior to analysis. Dimensionality reduction was performed using principal component analysis (PCA), and batch effects across independent ports were corrected with the harmony algorithm ^46^.

Clustering, doublet removal and cell type identification

Cells were clustered by constructing a shared nearest neighbour (SNN) graph and applying the Louvain community detection algorithm to identify distinct cell populations. The optimal number of principal components for dimensionality reduction was determined using an Elbow plot, and clustering resolution parameters were selected based on the endothelial cell complexity observed in each organ. Putative doublets were identified as cell clusters expressing high total UMI/gene counts, unusual gene co-expression and located in PCA outliers. Cells classified as singlets were retained and re-clustered. Final cluster identities were manually curated based on known endothelial cell subtypes and literature-based markers. R scripts used in this analysis are available at https://github.com/RuiBenedito/Benedito_Lab/.

Quantification of exon deletion

To assess the rate of exon deletion in *Vegfr2^flox/flox^*samples, we computed the number of unique molecular identifier (UMI) reads aligning to each exon, including the floxed region and the terminal 3′ exon/UTR (which is typically overrepresented in 10x Genomics 3′ libraries). Only reads with valid barcodes, UMIs, and matching strand orientation were retained.

Read counts for floxed exons were normalized to total UMI counts per cell. These values were added to the Seurat metadata. Deletion efficiency was visualized using dot plots and violin plots, comparing expression of the whole transcript (including the 3′ UTR).

Calculation of *sVegfr1/flVegfr1* ratio

To obtain the ratio between the soluble (*sVegfr1*) and full-length (*flVegfr1*) isoforms, the normalized reads from exon 13b (last exon in the soluble form) were divided by the normalized reads of exon 30 (last exon in the full-length version).

### Statistics and reproducibility

Most numerical data shown in charts was first compiled and processed with Microsoft Excel 2019 and after analyzed and plotted with Graphpad Prism v10.1.0. All bar graphs show mean ± standard deviation. The experiments were repeated with independent animals, as stated in the source data file or figure legends. The comparisons between two sample groups with a Gaussian distribution were by unpaired two-tailed Student *t*-tests. The comparisons among more than two groups were done by one-way or 2-way analysis of variance followed by multiple comparison tests. Datapoints were analyzed and plotted with GraphPad Prism. No randomization or blinding was used, and the animals or tissues were selected for analysis based on their genotype, the detected Cre/FlpO-dependent recombination frequency and the quality of multiplex immunostaining. The sample sizes were chosen according to the observed statistical variation and published protocols.

---

**Extended Data Figure 1:**
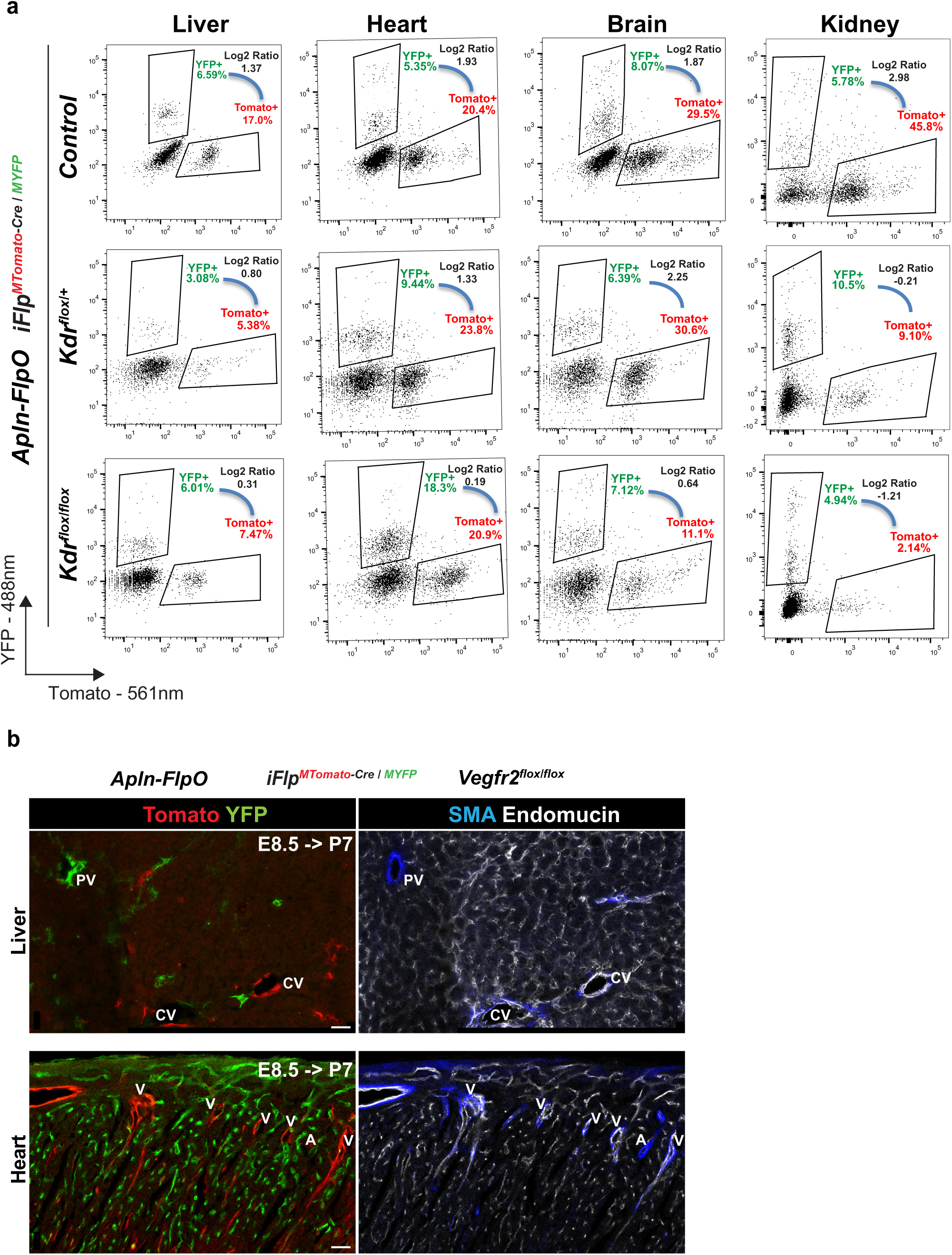
Ratiometric analysis of the expansion and mobilization of *Vegfr2^KO^* and wild-type cells. **a,** Representative FACs plots of CD31+ ECs collected from different organs (see quantitative data in Fig. 1b), showing YFP+ and Tomato+ cell frequencies, and their log2 ratio, in mice carrying the *iFlp^MTomato-Cre/MYFP^* allele in combination with the *Apln-FlpO* allele. **b,** Representative liver and heart confocal micrographs of *Apln-FlpO, iFlp^MTomato-Cre/MYFP^* -induced mosaic deletion of *Vegfr2* (*Vegfr2^KO-E8.5-P7^*), showing that Tomato+ *Vegfr2^KO^*ECs locate preferentially to venous vessels (Endomucin+ and SMA-). Scale bars are 25 μm. Abbreviations: PV-Portal vein, CV-Central vein, A – artery and V-vein.

**Extended Data Figure 2:**
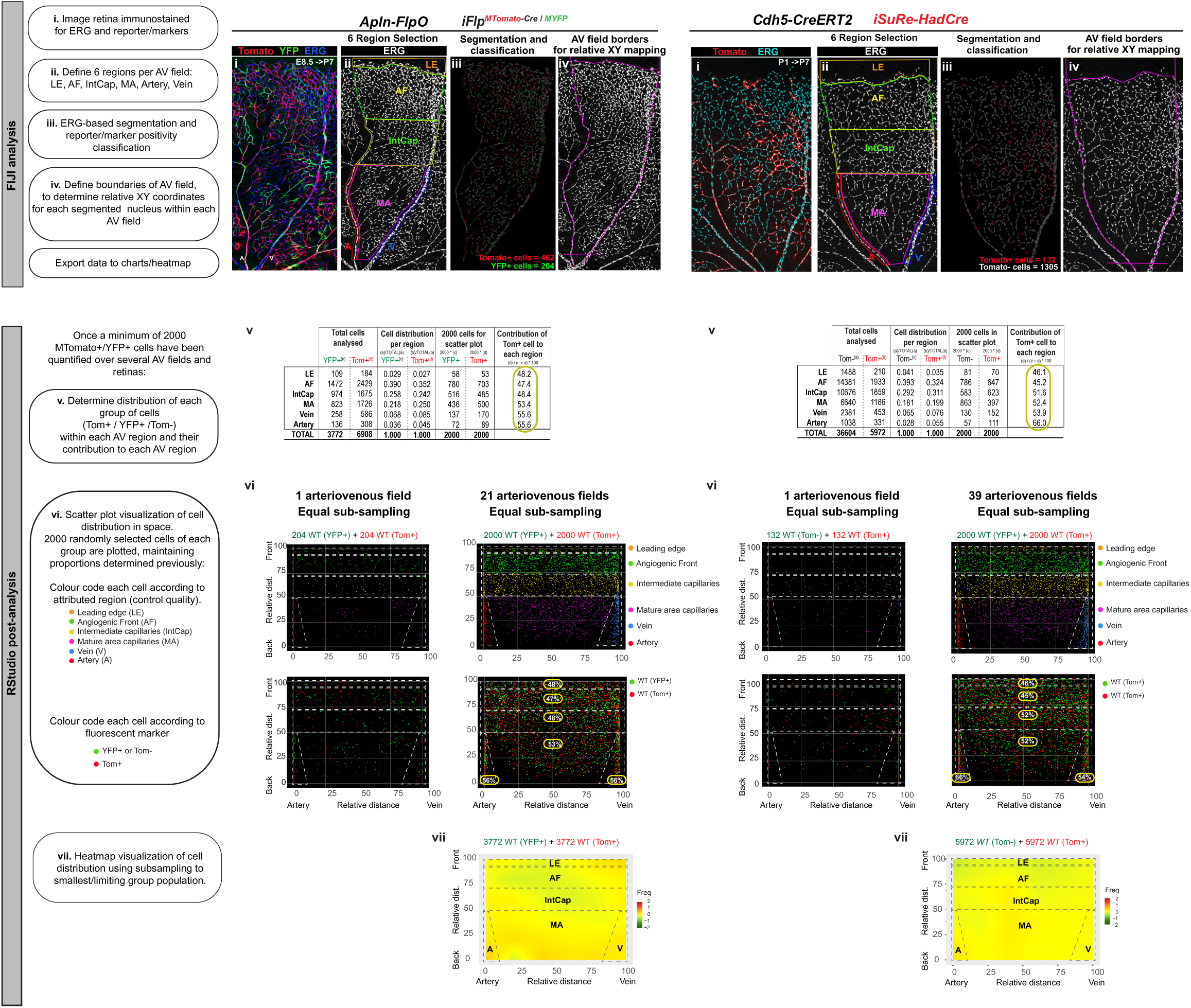
Cellular digital mapping and heatmap generation. Flow-chart depicting the main steps in image analysis used for the digital mapping of the identified cells, and the generation of heatmaps or XY dot plots. An example of the analysis for a control retinal AV field in an *AplnFlpO, iFlp^MTomato-Cre/MYFP^*or *Cdh5-CreERT2, iSure-HadCre* background are presented. Definition of the 6 regions of interest, ERG-based segmentation (endothelial nuclei), nuclear-classification according to reporter or marker expression and relative distance of each nucleus to the AV limits are determined using a custom made semi-automatic FIJI script (steps i-iv). Once a minimum of 2000 cells per reporter group (Tomato+/-, YFP+) are analysed, data is then post-analysed in RStudio to determine the cell distribution of each reporter group in each region and the contribution of Tomato+ cells to each region when compared to control (YFP+ or Tomato-cells) (step v). XY dot plots (step vi) and heatmaps (step vii) are then generated.

**Extended Data Figure 3:**
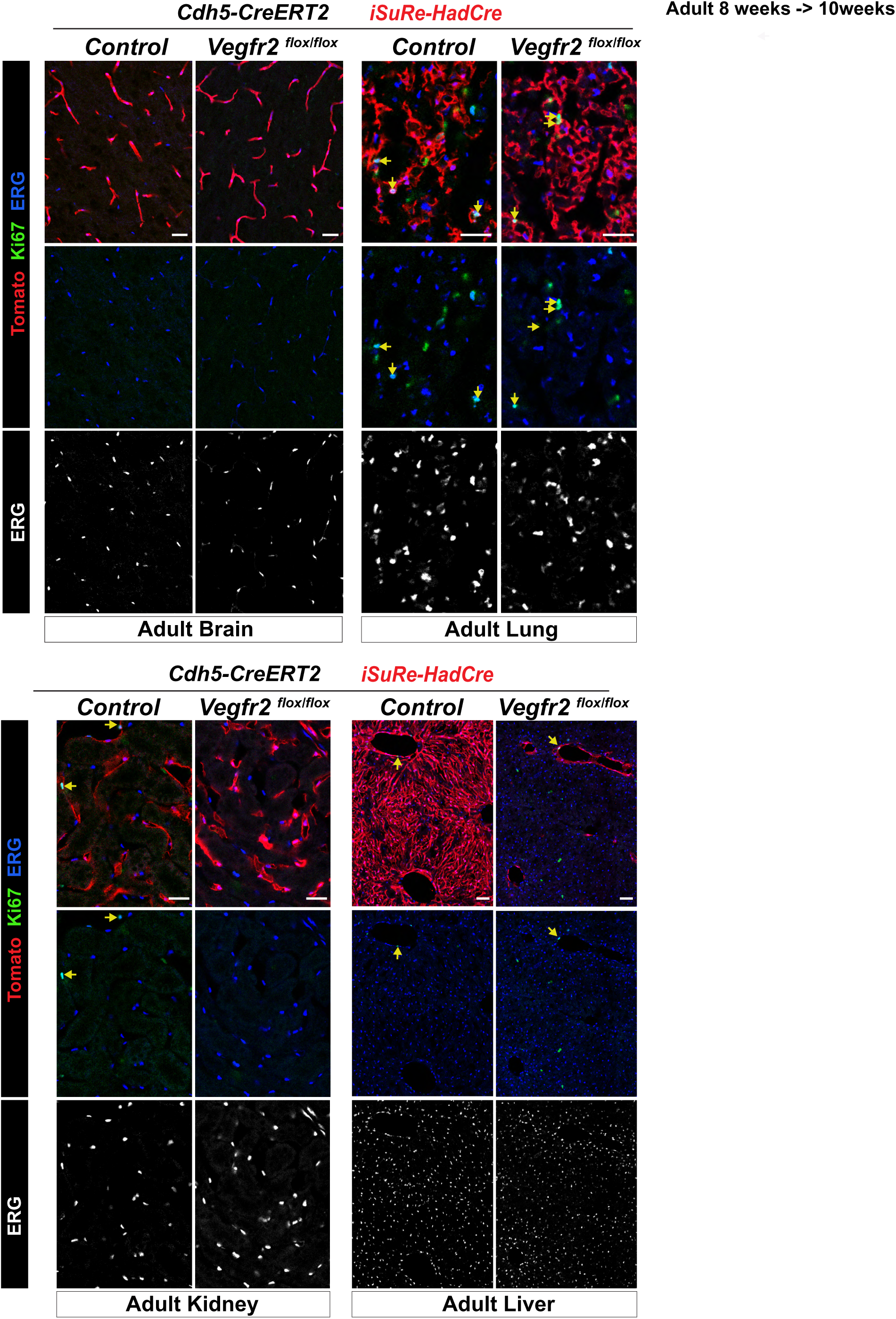
Impact of VEGFR2 deletion on the proliferation of adult ECs. Representative confocal micrographs of the indicated adult organs collected 2 weeks after induction with tamoxifen, and immunostained for Ki67 and ERG. Yellow arrows indicating Tomato+/Ki67+ ECs (see also Figure 3f for quantifications). Scale bars are 25μm.

**Extended Data Figure 4:**
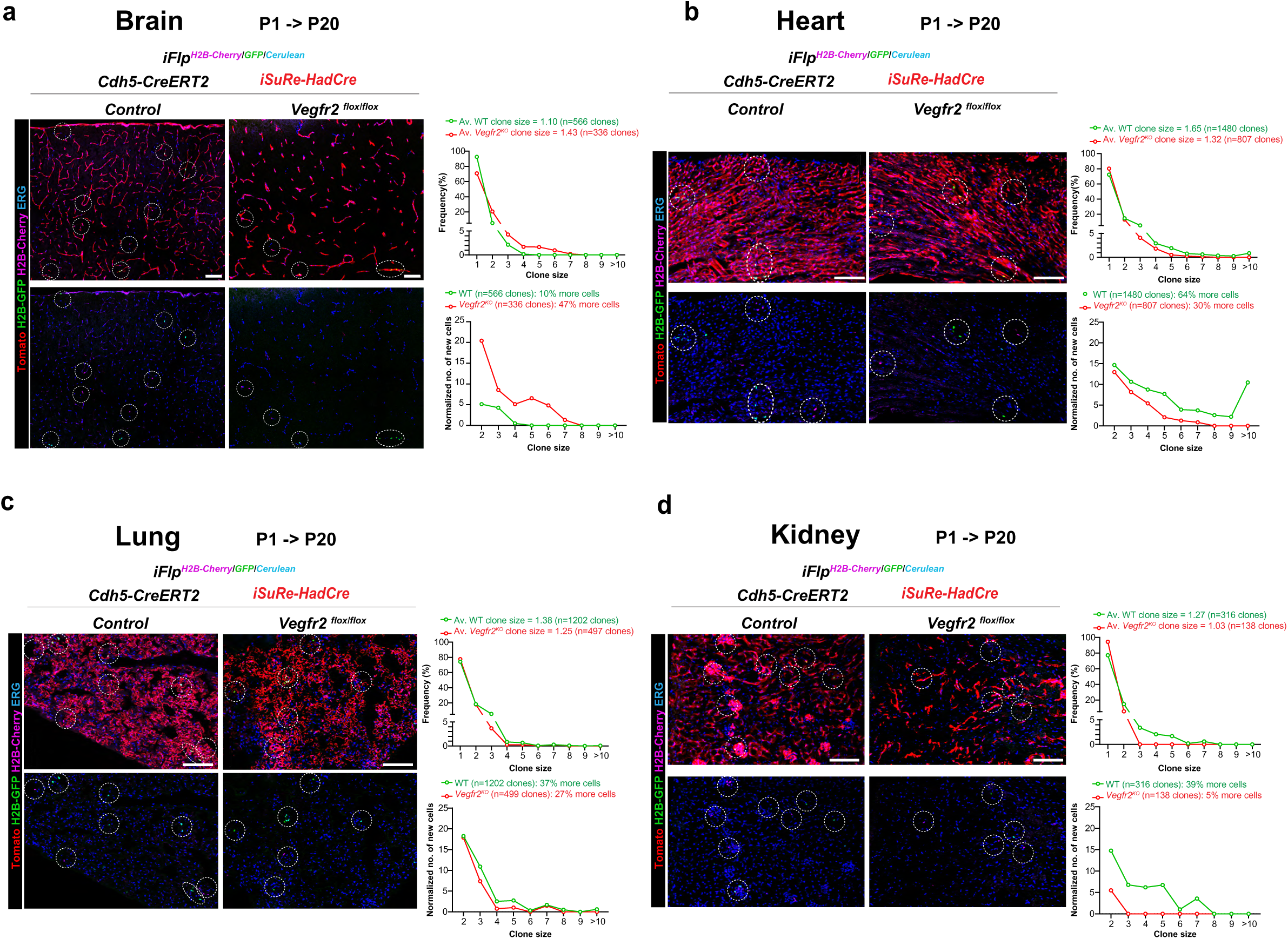
Clonal expansion analysis of control and *Vegfr2^KO^* ECs from P1 to P20. **a-d,** Representative confocal micrographs of the indicated organs from mice induced at P1 and collected at P20. Besides the *Cdh5-CreERT2* and the *iSure-HadCre* alleles, these animals have the *iFlp-Mosaic^H2B-Cherry/GFP/Cerulean^* reporter allele (Garcia-Gonzalez et al., 2025), that enables sparse and multispectral labelling of nuclei within a fraction of the Tomato+/mutant cells, giving scClonal resolution. Charts with the quantification of clone size and its frequency. Individual cells that have not expanded, or clones, are depicted by dotted circles. Scale bars are 100 μm.

**Extended data Figure 5.**
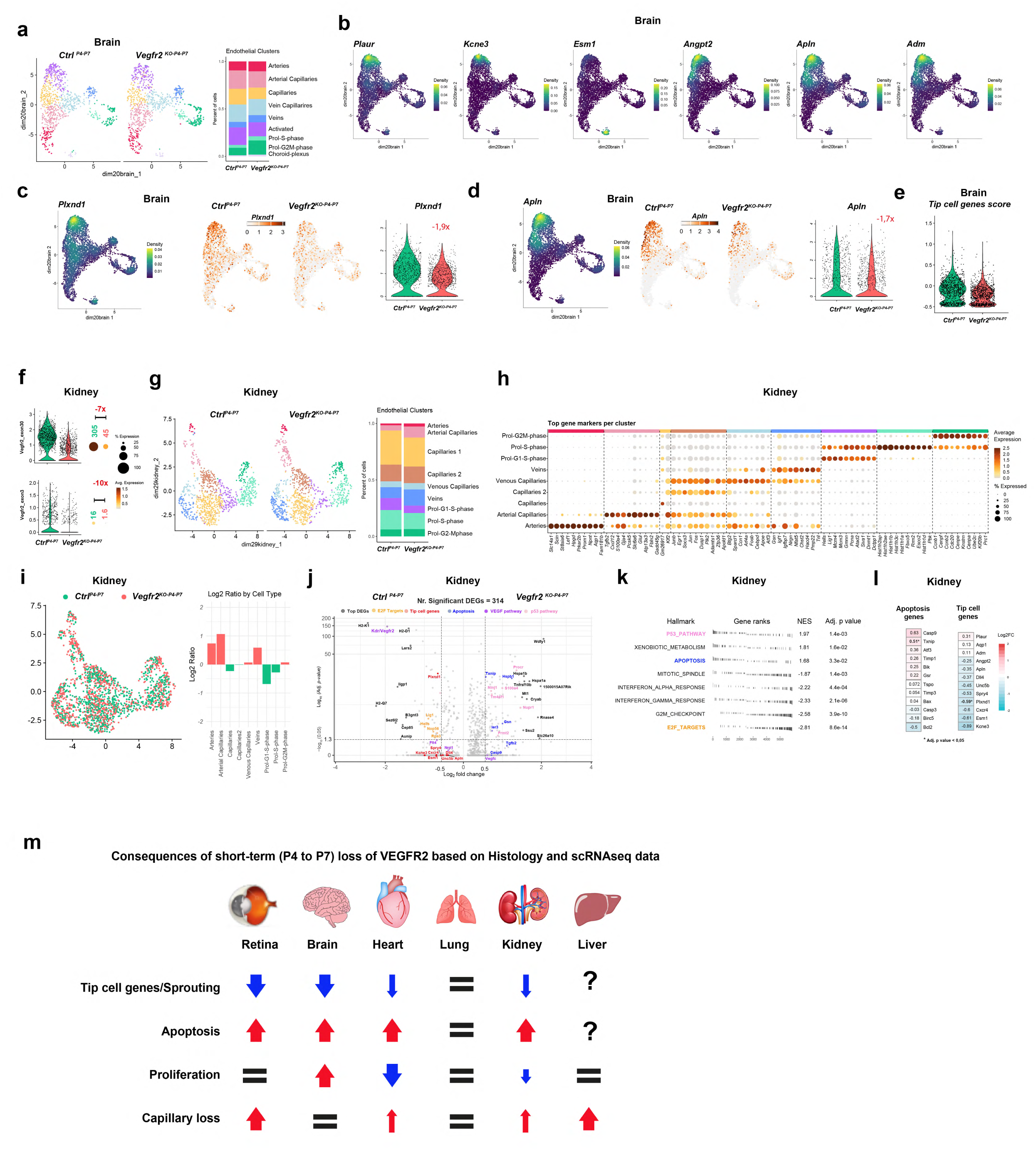
scRNAseq analysis of *Vegfr2^KO-P4-P7^* ECs. a,b, UMAP showing the different clusters. (a) and the expression of canonical tip cell marker genes in the activated cell cluster (b). **c,d**, Umaps and violin plots showing the expression of two genes known to be highly expressed by sprouting tip cells, and downregulated after *Vegfr2* deletion. **e**, Violins showing a tip cell genes score based on the combined expression of several tip cell genes (PlexinD1, Kcne3, Esm1). **f**, Violin plots showing expression of the terminal exon 30 (detected in the 3’10x genomics library sequencing) and the deleted floxed exon 3, used to assess *Vegfr2* deletion efficiency from P4 to P7. Note that in some cells there is still some residual *Vegfr2* mRNA given the short-term deletion nature of the assay, or some contamination with mRNA from wild-type kidney endothelial cells. Red negative values indicate fold changes between conditions. Green and red labels above the dots show the average expression multiplied by the percentage of expressing cells, representing absolute expression levels per group. **g**, UMAP and accompanying histobar plot illustrating the distribution and proportion of kidney endothelial cell clusters within each condition. **h**, Dot plot showing the expression of key marker genes for the different clusters. **i**, UMAP projection of the control and *Vegfr2^KO-P4-P7^* Kidney ECs. The associated bar plot shows the log2 ratio of the number of *Vegfr2^KO-P4-P7^* divided by the number of control cells per cluster. **j**, Volcano plot of showing top DEGs. Colored genes correspond to highlighted pathways or processes of biological relevance. **k**, List of hallmark pathways deregulated in kidney *Vegfr2^KO-P4-P7^* ECs. **l**, Heatmap showing the average log2 fold change of apoptosis-related and tip-cell genes in kidney endothelial cells between *Vegfr2*KO (P4–P7) and control samples.

**Extended data Figure 6.**
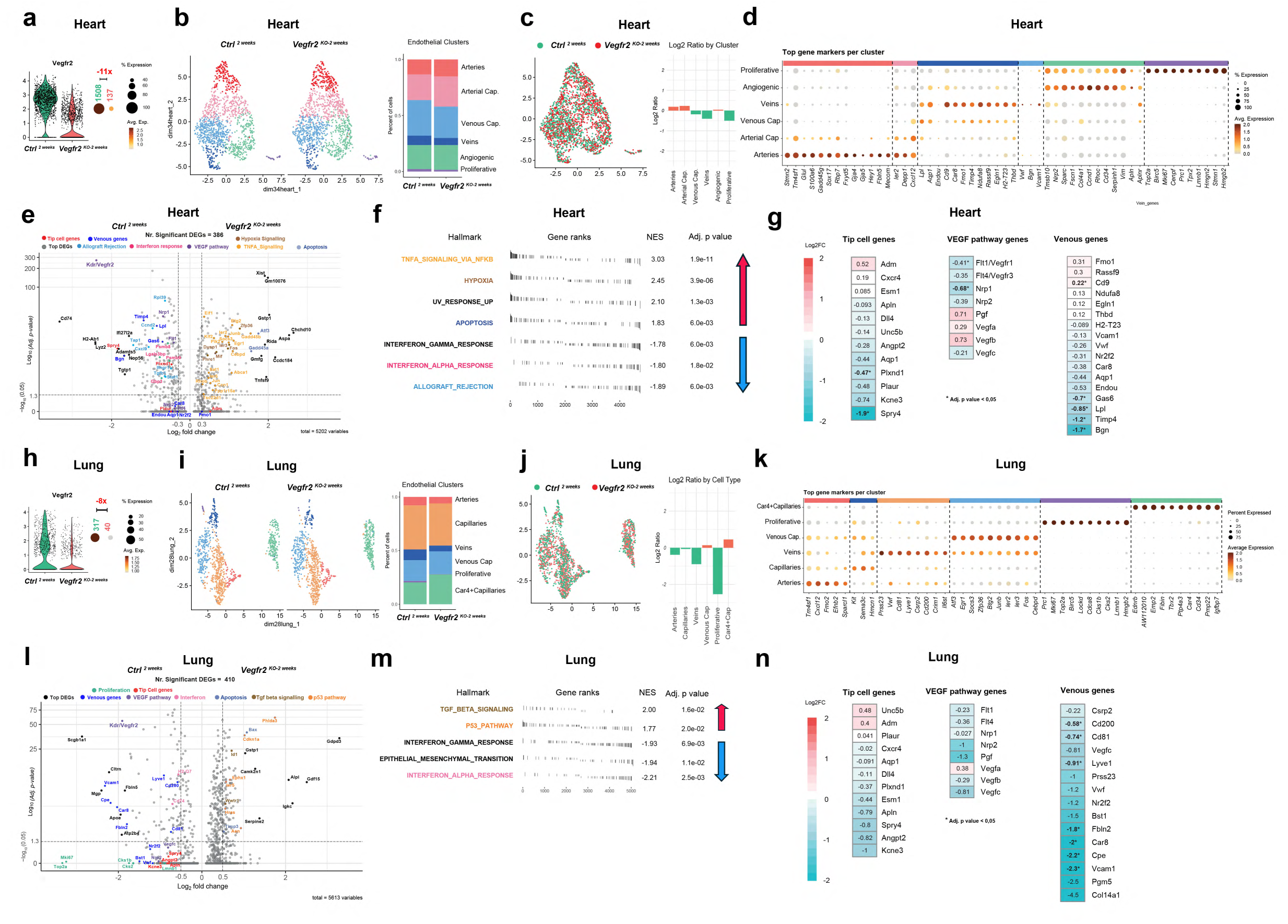
scRNAseq analysis of *Vegfr2^KO-2weeks^* ECs from the adult heart and lung. **a-g** Adult heart ECs scRNAseq data**. h-n,** Adult Lung ECs scRNAseq data. **a**, Violin plots showing expression of the whole *Vegfr2* mRNA. Red negative value indicate fold change between conditions. Green and red labels above the dots show the average expression multiplied by the percentage of expressing cells, representing absolute expression levels per group. **b,i** UMAP showing the identified EC clusters and histobar plots showing their relative proportions per condition. **c, j,** UMAP showing the spatial distribution of control and *Vegfr2^KO-2weeks^* ECs. The bar plot shows the log2 ratios of *Vegfr2^KO-2weeks^* ECs over control cells per EC cluster. **d, k,** Dot plot showing the top cluster marker genes. **e, l,** Volcano plot depicting the top DEGs and also other biologically significant genes when contrasting control and *Vegfr2^KO-2^ ^weeks^* ECs. **f, m,** Top significantly up and downregulated hallmark pathways in *Vegfr2^KO-2^ ^weeks^* ECs when compared to control ECs. Highlighted hallmarks match colour code used in e, l. **g, n,** Heatmap showing the average Log2FC of the indicated genes in *Vegfr2^KO-2^ ^weeks^* ECs when compared to control ECs.

**Extended data Figure 7.**
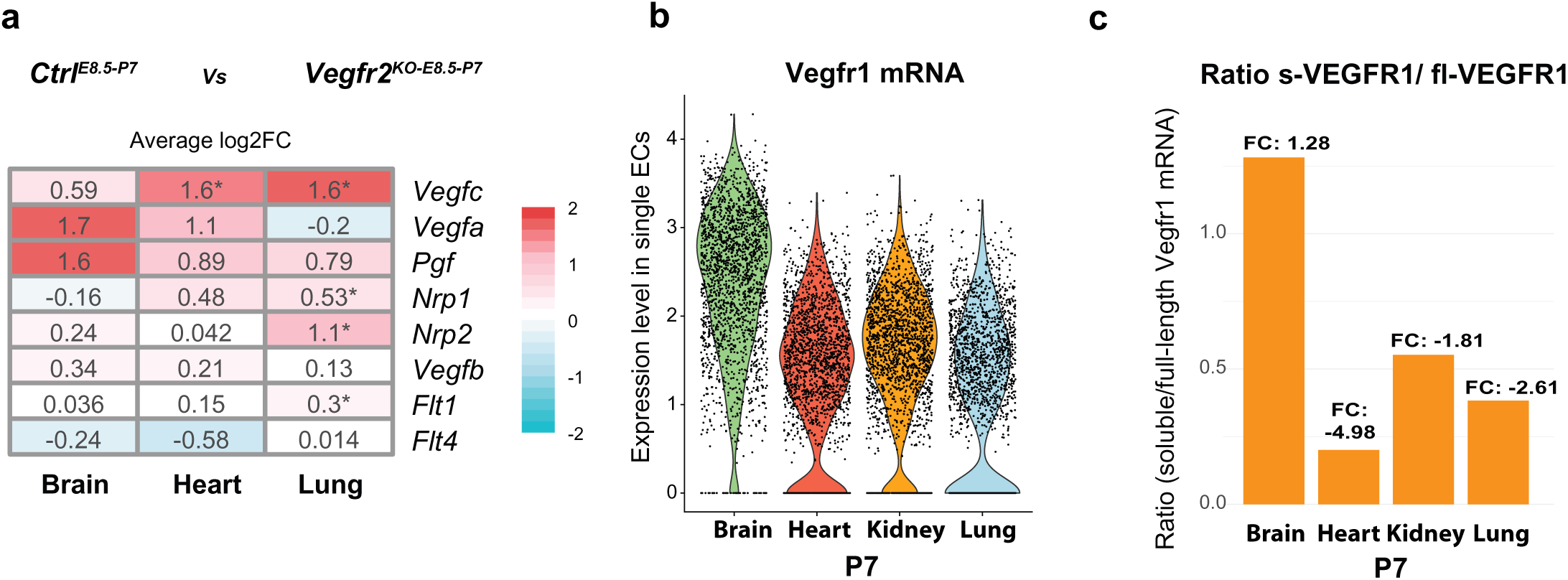
Expression of VEGF ligands and receptors in distinct postnatal organs ECs. **a,** Average log2FC expression in postnatal day 7 (P7) ECs for distinct VEGF pathway ligands and receptors. **b**, Violin plots showing expression of *Vegfr1* in distinct organs ECs. **c**, Ratio of soluble *Vegfr1* over full-length/membrane *Vegfr1* mRNA expression in distinct organs ECs.

